# Overcoming Preferred Orientation in Cryo-EM With Ultrasonic Excitation During Vitrification

**DOI:** 10.1101/2025.09.14.676144

**Authors:** Harry M. Williams, Wyatt A. Curtis, Michal Haubner, Jakub Wenz, Marcel Drabbels, Ulrich J. Lorenz

## Abstract

Preferred particle orientation remains a frequently encountered problem in cryo-electron microscopy that arises when proteins adsorb to the air-water interface in only a limited number of orientations. This issue can significantly increase the data acquisition time required to reach a desired resolution or even make it impossible to obtain a reconstruction altogether. Here, we show that preferred orientation can be overcome by continuously exciting the sample with ultrasonic waves during vitrification. Our experiments suggest that mechanical oscillations induced in the sample support continuously shake proteins loose from the air-water interface, thereby scrambling their orientations. The simple, physical nature of this mechanism should make it applicable to a wide range of proteins. Since our method can be easily implemented in existing vitrification devices, we expect it to find widespread adoption.

---

Significant efforts are being devoted to improving the reliability of sample preparation in cryo-electron microscopy (cryo-EM).^1–3^ Preferred particle orientation presents a frequently encountered issue that arises when proteins adsorb to the air-water interface with the hydrophobic parts of their surface, so that after vitrification, the cryo-EM sample contains only a limited number of viewing directions.^4,5^ In severe cases, this may limit the resolution of a single-particle reconstructions along some viewing directions or even make it impossible to obtain an interpretable map.^6^ While a range of methods have been developed to address preferred orientation, a simple, general solution has remained elusive. Tilting the specimen stage provides additional particle views, but also increases the ice thickness in the viewing direction, making this approach less suitable for small proteins.^7^ Adsorption to the air-water interface can also be reduced by chemically altering the interfacial interactions, for example by adding surface-active compounds^4,8^ or changing the type of specimen support film.^9,10^ However, such strategies require re-optimisation of the sample and are therefore time-consuming. Specialized sample preparation procedures can be used to reduce the time between sample application and vitrification, so that fewer proteins are able to reach the interface and adsorb there.^11–13^ However, conventional approaches are frequently not fast enough to outrun this process entirely.

We have recently shown that flash melting and revitrifying cryo-EM samples with microsecond laser pulses reduces preferred orientation.^14–16^ However, this method appears to be less effective for small proteins. For example, we were not able to improve the angular distribution of hemagglutinin, a 170 kDa protein that is notorious for its strong preferred orientation.^7,13,17^ Our experiments suggest that the laser pulse excites oscillations of the specimen support,^18,19^ so that the accompanying motions of the thin liquid film detach particles from the air-water interface and scramble their orientations. However, in a competing process, particles can also diffuse back to the interface and readsorb. This is particularly efficient for small proteins because of their short diffusion times, making it more difficult to improve their angular distributions. Here, we show that preferred orientation can be reduced for proteins of a range of sizes and symmetries by continuously exciting the sample with ultrasonic waves during vitrification. This notably allows us to overcome preferred orientation even for such a challenging case as hemagglutinin.

Figure 1 illustrates the experimental concept. Cryo-EM samples are prepared through jet vitrification^2^ while continuously exciting mechanical oscillations of the specimen support with an ultrasonic transducer (Fig. 1a). The motions induced in the thin liquid film continuously detach particles from the air-water interface and reshuffle their orientations, so that after vitrification, an improved angular distribution is obtained (Fig. 1b). Experiments are performed with a custom jet vitrification setup (Supplementary Information 1). Samples are prepared by applying 3.5 µL protein solution onto a clipped holey carbon specimen grid (1.2 µm holes, 1.3 µm apart on 200 mesh copper), and excess liquid is removed through single-sided blotting. The sample is then placed in the jet vitrification device, where it is excited with ultrasonic waves by a transducer located at a distance of 9 mm (SECO SC049, driven with a 490 kHz square wave of 100 V amplitude). After about 10 ms of ultrasonic excitation, the sample is then vitrified with a jet of liquid ethane. Finally, the specimen grid is transferred to a high-resolution electron microscope for imaging (Supplementary Information 2–5).

**Figure 1.**
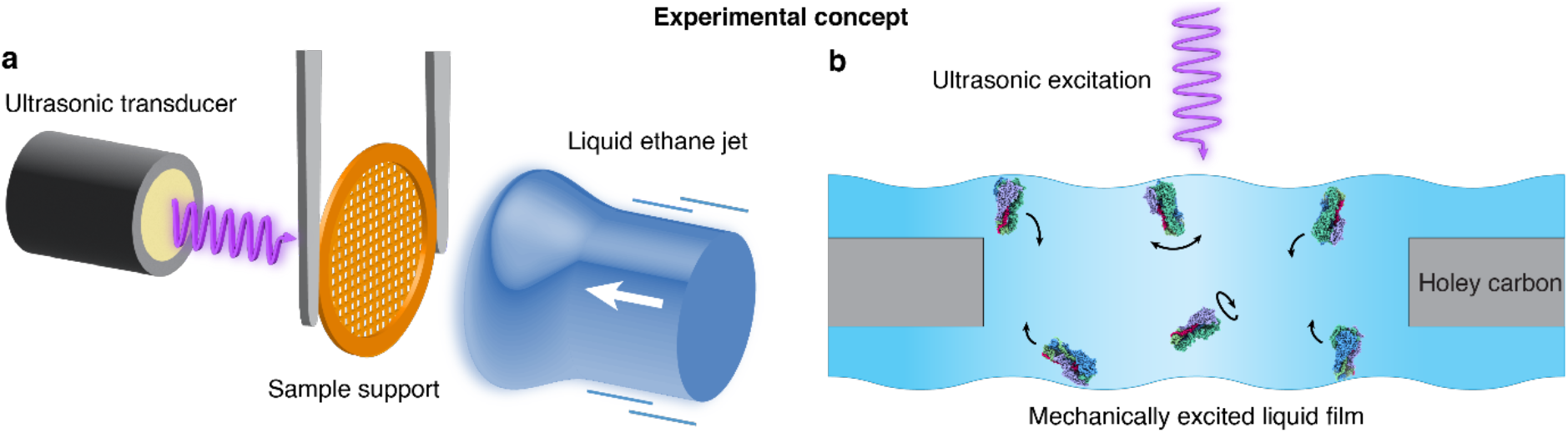
Experimental concept. **a** A cryo-EM sample is prepared through jet vitrification while mechanical oscillations of the specimen support are excited with an ultrasonic transducer. **b** The motions induced in the liquid film cause the particles to detach from the air-water interface and rotate freely. Vitrification therefore traps the particles in a non-equilibrium angular distribution, yielding a sample in which preferred orientation is reduced.

Figure 2a demonstrates that ultrasonic excitation reduces preferred orientation for a sample of the 50S ribosomal subunit (1.34 MDa, point group C_1_). A reconstruction of a conventional, jet vitrified sample is shown in the top together with the corresponding angular distribution of the particles, which reveals several pronounced maxima, indicating strong preferred orientation. If the sample is prepared with ultrasonic excitation, but under otherwise identical conditions (bottom), the angular distribution noticeably broadens, and additional maxima appear. The sampling compensation factor (SCF*), a measure of the isotropy of the angular distribution, improves from 0.66 to 0.83. The SCF* can take values between 0 and 1, where 1 describes a sample in which all viewing directions are equally represented. Because of the more homogeneous angular distribution, the resolution of the reconstruction improves from 2.8 Å to 2.6 Å. Moreover, the streaky artefacts are noticeably reduced that are visible in the map obtained from the conventional sample and that are a consequence of preferred orientation (Supplementary Information 2).

**Figure 2.**
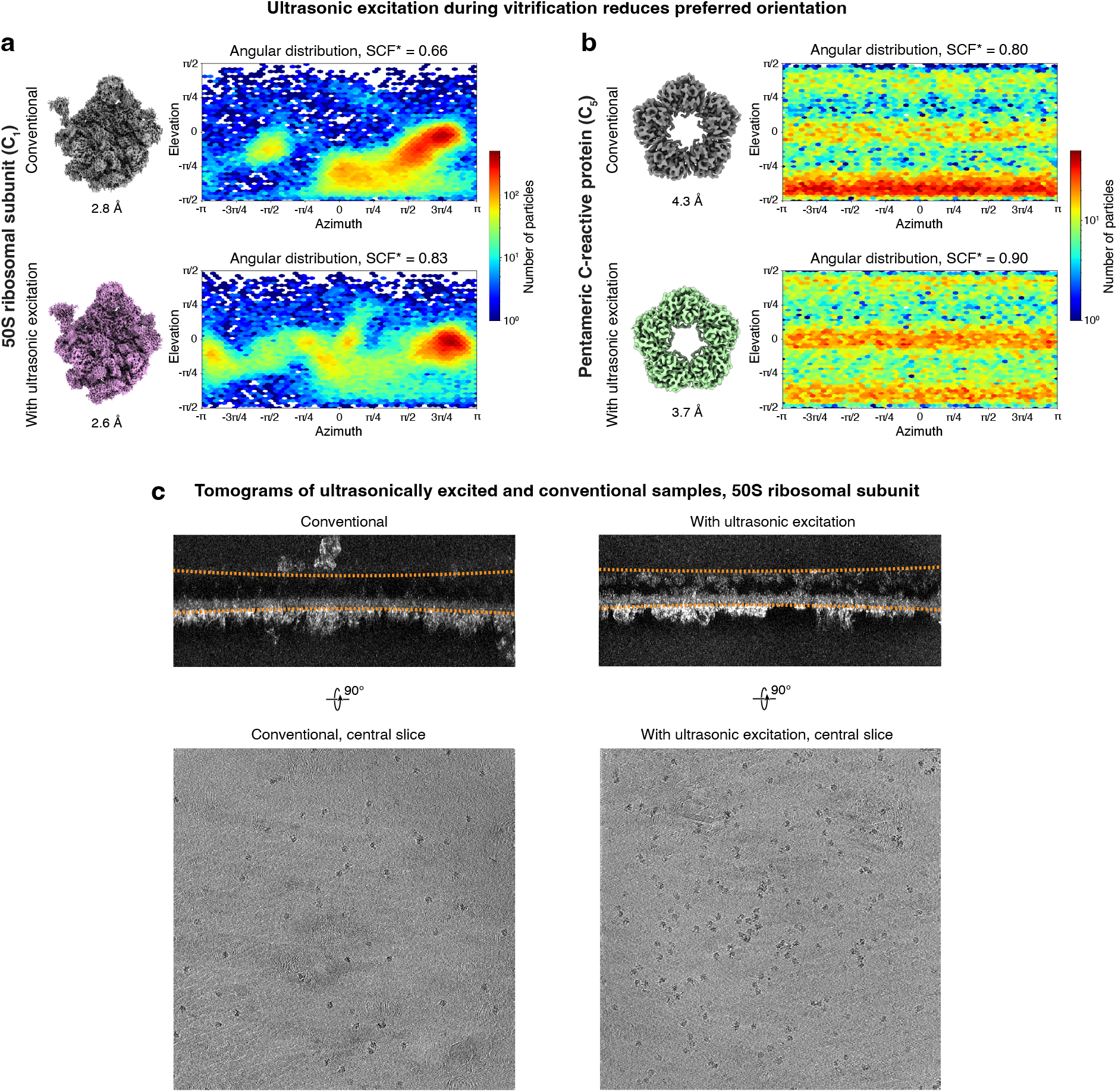
Ultrasonic excitation during vitrification reduces preferred orientation. **a**,**b** Reconstructions and angular distributions from conventional, jet vitrified samples as well as samples that were excited with ultrasonic waves during jet vitrification for the 50S ribosomal subunit (**a**) and the pentameric C-reactive protein (**b**). With ultrasonic excitation, both the sampling compensation factor (SCF*), a measure of the isotropy of the angular distribution, as well as the resolution of the reconstructions improve. **c** Tomograms of conventional and ultrasonically excited samples of the 50S ribosomal subunit (the sample interface is indicated with a dashed line). A slice through the center of the sample reveals that ultrasonic excitation causes a larger number of particles to detach from the air-water interface.

Ultrasonic excitation during jet vitrification similarly reduces preferred orientation for the pentameric form of human C-reactive protein (120 kDa, C_5_, Fig. 2b). The SCF* improves from 0.80 to 0.90, and the resolution from 4.3 Å to 3.7 Å. At the same time, the quality of the map increases significantly (Supplementary Information 4). Note that it was previously reported that the C-reactive protein pentamer and decamer have such strong preferred orientation that it is not possible to obtain a reconstruction without using surfactants.^20,21^ Here, we are able to do so even without ultrasonic excitation (Supplementary Information 4), which is potentially due to the different vitrification method we employ. Tomograms of the 50S ribosomal subunit samples provide further insights into how ultrasonic excitation alters the particle distribution (Fig. 2c). In a conventional sample, most particles are attached to the bottom interface, while under ultrasonic excitation, both interfaces are populated more evenly. Notably, slices taken through the center of the sample reveal a larger number of particles that are not attached to either interface.

We demonstrate that ultrasonic excitation can overcome preferred orientation even for hemagglutinin (170 kDa, C_3_), which is notorious for its strong preferred orientation.^7,13,17^ Figure 3a shows that we can only obtain a low-quality reconstruction from a conventional sample, with the bottom half of the protein density missing (2.7 Å nominal resolution).^14^ This is because the angular distribution is strongly dominated by top and bottom views (SCF* 0.25). In contrast, exciting the sample with ultrasonic waves during jet vitrification allows us to obtain a complete map of the protein (Fig. 3b). At a resolution of 2.7 Å, side chain densities are clearly visible. The corresponding angular distribution (SCF* 0.81) features three additional, symmetry equivalent maxima, revealing that ultrasonic excitation populates side views that are practically absent in a conventional sample (2D class averages in Fig. S12). Note that the fraction of side views in the sample is likely lower than suggested by the angular distribution, since the heterogeneous refinement steps that yield the reconstruction in Fig. 3b remove many top and bottom views (Supplementary Information 5–6). If we instead try to estimate the angular distribution in the sample by refining all particles against the volume in Fig. 3b, we obtain SCF* values of 0.43 and 0.56 for the conventional and ultrasonically excited sample, respectively (Supplementary Information 6).

**Figure 3.**
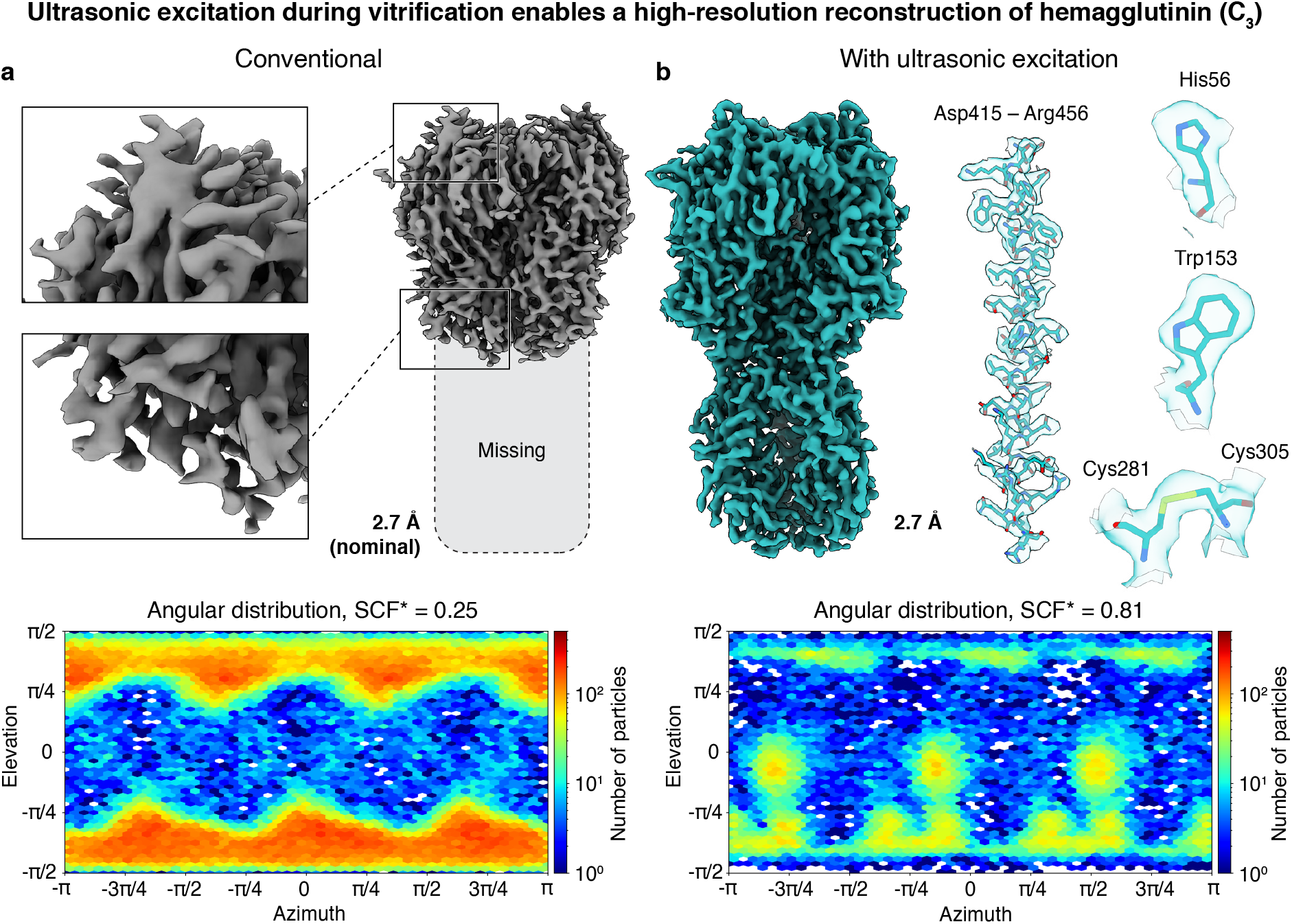
Ultrasonic excitation during vitrification enables a high-resolution reconstruction of hemagglutinin, a notoriously challenging case of preferred orientation. **a**,**b** Reconstructions and angular distributions from a conventional, jet vitrified sample as well as a sample that was excited with ultrasonic waves during jet vitrification. Because of strong preferred orientation, the conventional sample only yields a reconstruction in which the bottom half of the density is missing (**a**). In contrast, ultrasonic excitation improves the angular distribution by populating side views and restores the missing density (**b**). Details of the reconstruction are shown with a model jiggle-fitted into the map (PDB ID: 7VDF^22^). Note that if we estimate the angular distributions of both samples from a refinement of all particles against the volume in (**b**), we obtain SCF* values of 0.43 of 0.56 for the conventional and ultrasonically excited sample, respectively (Supplementary Information 6).

Our experiments suggest a simple mechanism by which ultrasonic excitation reduces preferred particle orientation. The ultrasonic waves likely excite flexural modes of the freestanding specimen support film, which has estimated eigenfrequencies of tens to hundreds of kilohertz, in a range close to the 490 kHz excitation frequency.^23^ The accompanying motions of the liquid film exert small forces on the proteins that detach them from the air-water interface and scramble their orientations. At the same time, particles diffuse back to the sample surface and readsorb in their preferred orientation. We counteract this competing process by continuously exciting oscillations of the specimen support, so that proteins are constantly being detached from the air-water interface. Under these conditions, the sample settles into a dynamic equilibrium, in which the fraction of particles adsorbed in their preferred orientation is reduced.

Interestingly, ultrasonic excitation leads to the appearance of additional maxima in the angular distribution of hemagglutinin, which evidently arise from particles that are adsorbed to the air-water interface in side views (Fig. 3). We estimate that their population is increased by about one order of magnitude relative to the fraction of particles adsorbed in top and bottom views. This suggests that ultrasonic excitation alters the relative rates with which particles with different orientations detach from the interface and readsorb. It is conceivable that particles adsorbed in top and bottom views experience a greater torque when exposed to shear motions of the liquid film and therefore detach more readily compared to particles adsorbed with their sides. It is also possible that the attachment rate of the top and bottom views decreases relative to the rate of the side views. Either effect would increase the population of the side views in the dynamic equilibrium.

In conclusion, ultrasonic excitation of the sample during vitrification provides a straightforward method for overcoming preferred particle orientation, while leaving the proteins intact, as confirmed by comparison to available molecular models (Fig. 3b and Fig. S9). For unmodified, bare hemagglutinin, which is notorious for its strong preferred orientation, our approach yields one of the highest-resolution structures reported to date, with only one other study achieving a marginally higher resolution, where surfactants were used to reduce adsorption to the air-water interface.^22^ This however resulted in a significantly worse B factor of 185.6 Å^2^, compared to 66.1 Å^2^ in our work. Since our method employs a simple, physical process for detaching proteins from the air-water interface and reshuffling their orientations, it should be applicable to a wide range of proteins. Likely, preferred orientation can be reduced even further by increasing the amplitude of the oscillations induced in the specimen support, either by matching the excitation frequency more closely to one of the eigenfrequencies of the sample or by increasing the power delivered by the ultrasonic transducer. Finally, it should be straightforward to implement our method in existing vitrification devices. For example, we present a proof-of-concept implementation for a plunge freezing device in Supplementary Information 7, where the sample moves past several ultrasonic transducers during the plunging process. This allow us to obtain an even better angular distribution for hemagglutinin (with the same method as above, we estimate the SCF* of the entire sample to be 0.74, compared with 0.56 in our jet vitrification experiment), allowing us to improve the resolution of the reconstruction to 2.0 Å and the B factor to 48.1 Å^2^. We note that instead of exciting oscillations of the sample through air, it should also be possible to transmit the ultrasonic waves more efficiently through the fixture holding the specimen grid.

## Acknowledgements

The authors would like to thank the EPFL ISIC electrical and mechanical workshops for their help in realizing this project, in particular S. Dutoit, B.C. Le Geyt, and G. Pasche, as well as B. Beckert from the Dubochet Center for Imaging Lausanne for providing a sample of the 50S ribosomal subunit. Cryo-EM and cryo-ET data collection was performed at the Dubochet Center for Imaging Lausanne (a joint initiative from EPFL, UNIGE, UNIL, UNIBE) with the assistance of A. Myasnikov, B. Beckert, S. Nazarov, I. Mohammed, and E. Uchikawa. The work was supported by Swiss National Science Foundation Grant TMCG-2_213773 (UJL) and by the Duke Center for HIV Structural Biology, NIH grant U54AI170752 (UJL).

## Author contributions

Conceptualization: MH, WAC, HMW

Methodology: MH, WAC, HMW, JH

Investigation: MH, WAC, HMW, JH

Visualization: MH, WAC, HMW, JH

Funding acquisition: UJL

Project administration: UJL

Supervision: UJL, MD

Writing – original draft: MH, WAC, HMW, JH

Writing – review & editing: MH, WAC, HMW, JH, UJL, MD

## Data and materials availability

Supplementary Information is available for this paper.

The cryo-EM maps have been deposited in the Electron Microscopy Data Bank (EMDB) and the Electron Microscopy Public Image Archive (EMPIAR) under accession codes EMD-54830 and EMPIAR-12979 (50S conventional sample), EMD-54829 and EMPIAR-12978 (50S ultrasonically excited sample), EMD-54832 and EMPIAR-12981 (recombinant hemagglutinin trimer conventional sample), EMD-54831 and EMPIAR-12980 (recombinant hemagglutinin trimer ultrasonically excited sample), EMD-54834 and EMPIAR-12983 (human C-reactive protein conventional pentameric sample), EMD-54833 and EMPIAR-12982 (human C-reactive protein ultrasonically excited pentameric sample), EMD-54836 and EMPIAR-12983 (human C-reactive protein conventional decameric sample), EMD-54835 and EMPIAR-12982 (human C-reactive protein ultrasonically excited decameric sample), and EMD-58181 and EMPIAR-XXXXX (recombinant hemagglutinin trimer conventional plunged sample), EMD-58182 and EMPIAR-XXXXX (recombinant hemagglutinin trimer ultrasonically excited plunged sample).

