## Supplementary Information for "Overcoming Preferred Orientation in Cryo-EM With Ultrasonic Excitation During Vitrification"

**This PDF file includes:**

1 | Cryo-EM sample preparation

2 | Data processing and analysis – 50S ribosomal subunit

3 | Cryo-ET of a sample of the 50S ribosomal subunit

4 | Data processing and analysis – C-reactive protein

5 | Data processing and analysis – hemagglutinin

6 | Angular distribution in the conventional and ultrasonically excited hemagglutinin samples

7 | Proof-of-concept implementation of ultrasonic excitation in a plunge-freezing device

8 | Cryo-ET data collection statistics

9 | Cryo-EM data collection and processing statistics

10 | References

### 1 | Cryo-EM sample preparation

Cryo-EM samples are vitrified with a custom jet vitrification device that we integrated into a commercial plunge freezing apparatus (Thermo Fisher Vitrobot Mark IV) and that we have described in detail previously.[1] Grids are prepared in the climate chamber of the plunge freezing apparatus. After single-sided blotting, the sample is lowered into the jet vitrification unit, which is mounted underneath the climate chamber. The sample is then excited with ultrasonic waves and vitrified with a jet of liquid ethane, after which the grid is immediately lowered further into a liquid ethane filled container that is placed underneath the jet vitrification device. From there, it is then transferred to liquid nitrogen and subsequently to a high-resolution electron microscope for imaging.

A purified sample of the 50S ribosomal subunit (40 $OD_{260}$/mL, 20 mM HEPES buffer, pH 7.5, 100 mM NaCl, 2 mM $MgCl_2$) was provided by Dr. Bertrand Beckert of the Dubochet Center for Imaging in Lausanne. Purified hemagglutinin was purchased from MyBioSource (MBS434205) in lyophilized form and resuspended in 1x PBS, pH 7.5 to a final concentration of 0.75 mg/mL. C-reactive protein from human fluids was purchased from Sigma-Aldrich (C4603) and used as is (2.6 mg/mL, 20 mM Tris, 280 mM NaCl, 5 mM $CaCl_2$, pH 7.8–8.2, 0.1% $NaN_3$).

Samples for jet vitrification are prepared in the climate chamber of the plunge freezing apparatus by applying 3.5 µL of the protein solution onto holey carbon grids (R1.2/1.3, 200 mesh copper, Quantifoil) that have been rendered hydrophilic (air glow discharge for 90 s, EasiGlow, Ted Pella) and clipped prior to use. After single-sided blotting (20 °C, 95% relative humidity, blotting force 0, 2 s blotting time for 50S ribosomal subunit; 4 °C, 90% relative humidity, blotting force 10, 4.5 s blotting time for hemagglutinin; 20 °C, 95% relative humidity, blotting force 0, 3.5 s blotting time for the C-reactive protein), the sample is then lowered into the jet vitrification device.

Samples are vitrified with a jet of liquid ethane. The jet is created by condensing ethane gas into a brass reservoir cooled to 98 K and then expelling the cryogenic liquid (about 0.3 mL) through a tube (2 mm inner diameter) with a pulse of compressed nitrogen. The jet travels with a speed of about 2.6 m/s when it reaches the sample. The sample is then immediately lowered further into a liquid ethane filled container. To improve the angular distribution of the particles, the grids are mechanically excited with ultrasonic waves for about 10 ms prior to the arrival of the liquid ethane jet. To this end, an ultrasonic transducer (SECO SC049) is placed at a distance of 9 mm from the sample and is excited with a 490 kHz square wave of 100 V amplitude, providing a sound intensity level of about 126 dB at the sample location, as estimated based on the specifications of the manufacturer.

### 2 | Data processing and analysis – 50S ribosomal subunit

High-resolution micrographs of a conventional cryo sample of the 50S ribosomal subunit as well as a sample that was subjected to ultrasonic excitation were collected at the Dubochet Center for Imaging in Lausanne, using a Thermo Fisher Titan Krios G4i transmission electron microscope equipped with a Falcon IV direct electron detector. Ultrasonic excitation does not alter the quality of the jet-vitrified samples, as illustrated in Fig. S1, which shows an atlas of a typical sample together with representative micrographs of individual areas.

Single-particle reconstructions were performed using CryoSPARC v4.7.1[2], with the corresponding workflows illustrated in Figs. S2 and S3. The data acquisition parameters and processing statistics are summarized in Table S2. Micrographs were subjected to patch motion correction and patch CTF estimation. Micrographs with an estimated resolution of worse than 10 Å were rejected. Particles from the conventional and ultrasonically excited samples were picked using a blob picker with a diameter ranging from 150 to 400 Å. The particles were extracted with a box size of 784 px and Fourier-cropped to 196 px.

For the conventional dataset, particles were first subjected to three rounds of 2D classification followed by *ab initio* reconstruction (three classes, $C_1$ symmetry). Particles from the two best *ab initio* classes were homogeneously refined ($C_1$ symmetry) into a single volume. This volume and two decoys were then used in two rounds of heterogeneous refinement (three classes, $C_1$ symmetry). Particles from the best class resulting from the final round of heterogeneous refinement were re-extracted with a box size of 784 px. A random subset of 50,000 of these re-extracted particles were then homogeneously refined ($C_1$ symmetry) producing the final volume shown in Fig. S2.

For the ultrasonically excited dataset, particles were subjected to three rounds of 2D classification followed by *ab initio* reconstruction (three classes, $C_1$ symmetry). Particles from the best *ab initio* classes were homogeneously refined ($C_1$ symmetry) into a single volume. This volume and two decoys were then used in two rounds of heterogeneous refinement (three classes, $C_1$ symmetry). Particles from the best class resulting from the final round of heterogeneous refinement were re-extracted with a box size of 784 px. A random subset of 50,000 of these re-extracted particles were homogeneously refined ($C_1$ symmetry) producing the final volume shown in Fig. S3.

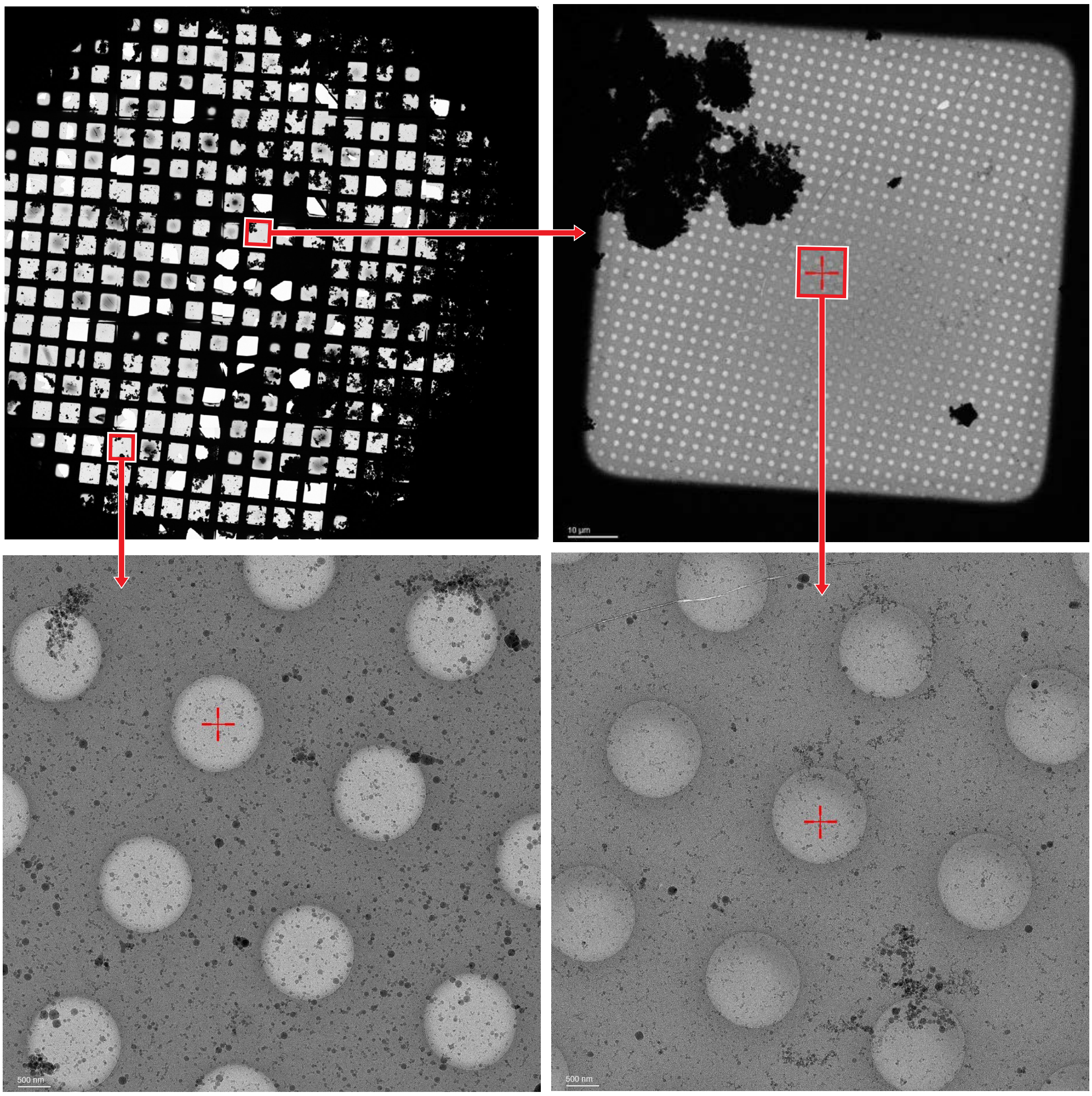


**Figure S1 | Atlas of a typical sample of the 50S ribosomal subunit prepared with ultrasonic excitation, together with representative micrographs of individual areas.** Ultrasonic excitation does not alter the quality of the jet-vitrified samples.

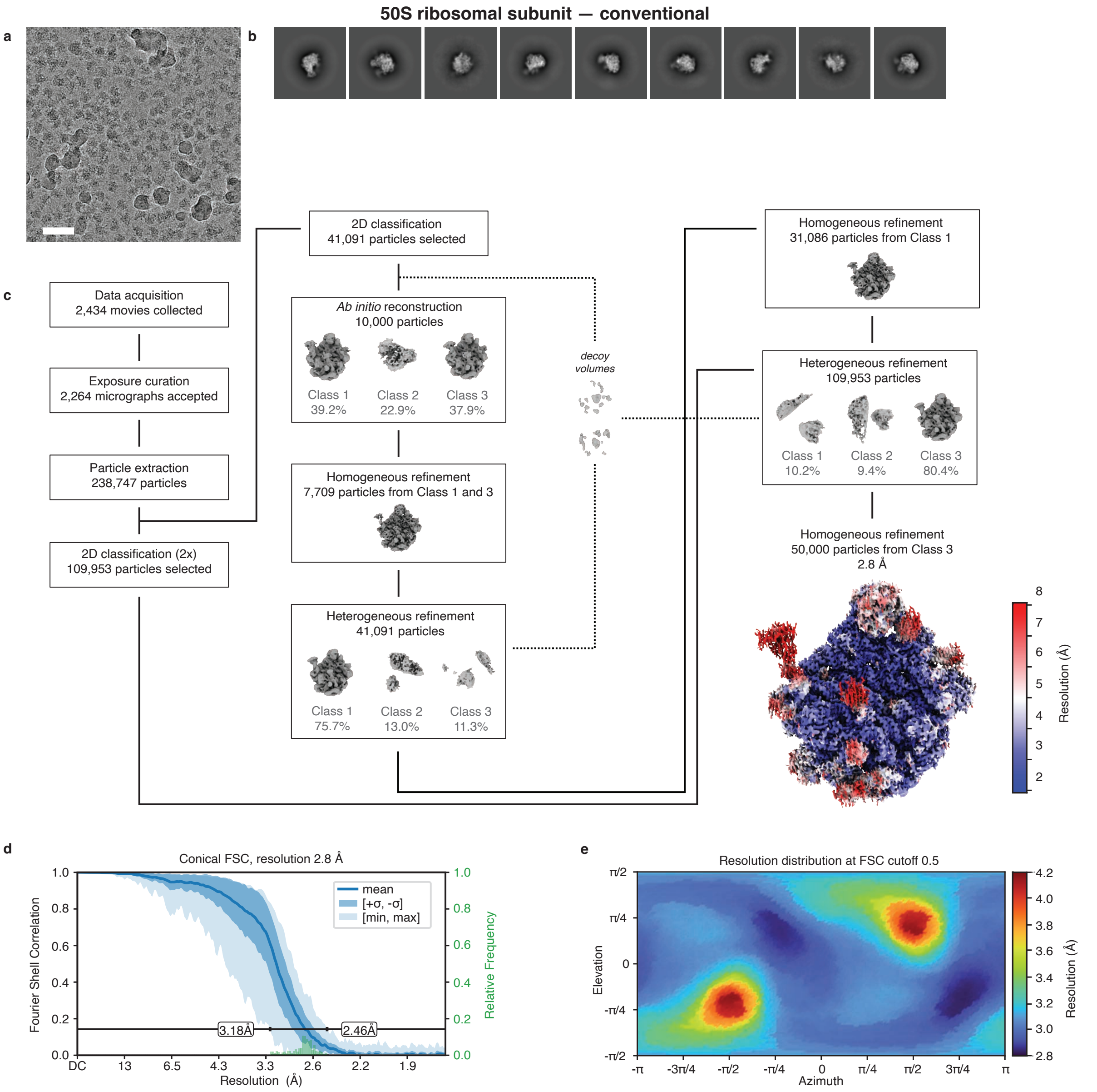


**Figure S2 | Processing workflow for the conventional sample of the 50S ribosomal subunit.** **a** Representative micrograph. Scale bar, 500 Å. **b** Selected 2D class averages. **c** Data processing workflow in CryoSPARC. All reconstructions are performed with $C_1$ symmetry. The volumes are displayed at a level of 5σ above the mean. The final map is displayed at a level 3σ above the mean, with the local resolution estimation at the FSC cutoff of 0.5 indicated in color. **d** Conical FSC, with the 0.143 threshold indicated by a black line. The mean FSC value is shown as a solid blue line. Dark blue shading indicates one standard deviation of the FSC curve, while the light blue shading represents the minimum and maximum FSC values. A histogram of the resolution values obtained from the conical FSC is shown in green. **e** Angular distribution of the resolution at FSC cutoff 0.5, generated using the 3DFSC file from Orientation Diagnostics job in CryoSPARC.

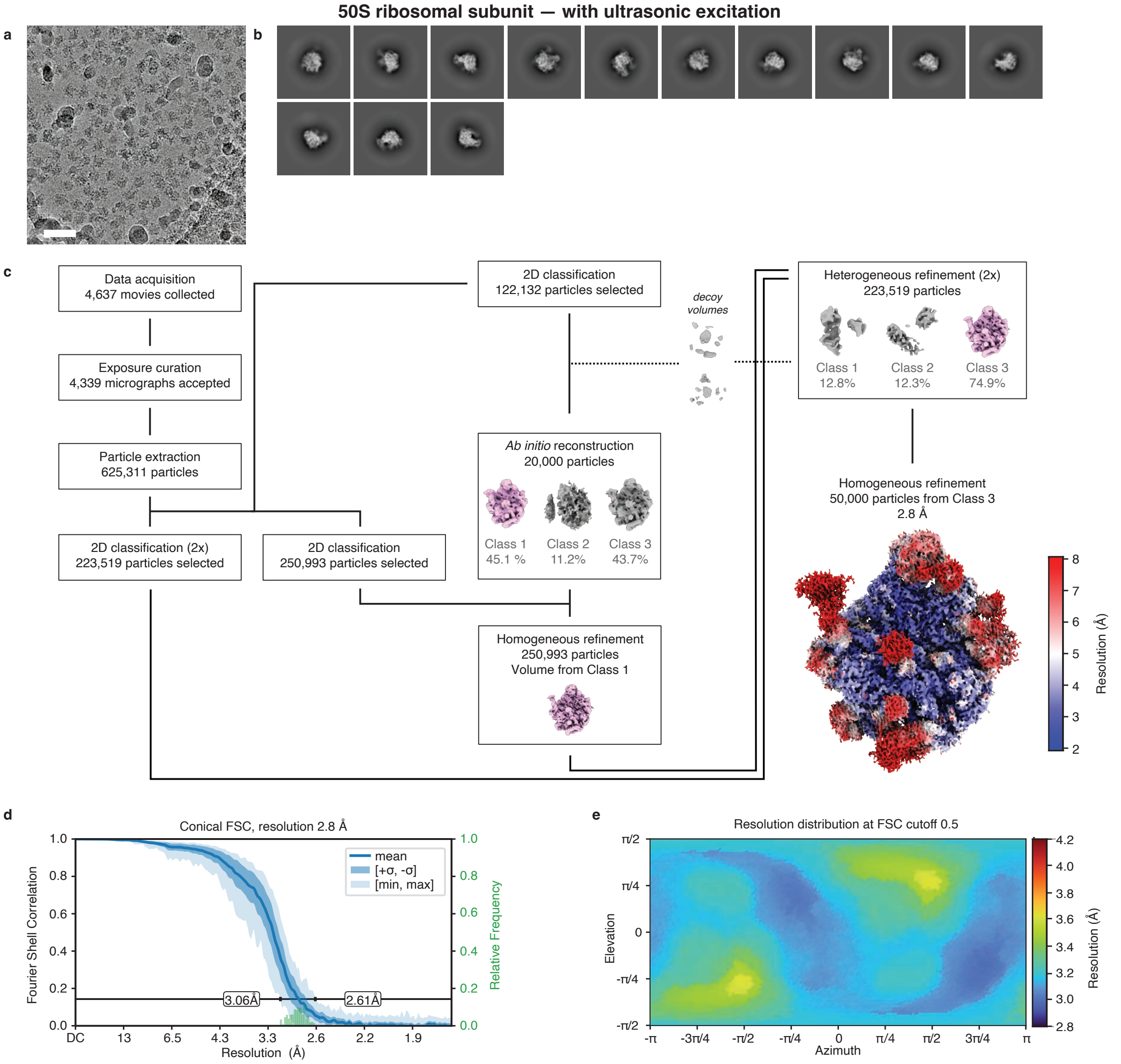


**Figure S3 | Processing workflow for the ultrasonically excited sample of the 50S ribosomal subunit. a** Representative micrograph. Scale bar, 500 Å. **b** Selected 2D class averages. **c** Data processing workflow in CryoSPARC. All reconstructions are performed with $C_1$ symmetry. The volumes are displayed at a level of 5σ above the mean. The final map is displayed at a level 3σ above the mean, with the local resolution estimation at the FSC cutoff of 0.5 indicated in color. **d** Conical FSC, with the 0.143 threshold indicated by a black line. The mean FSC value is shown as a solid blue line. Dark blue shading indicates one standard deviation of the FSC curve, while the light blue shading represents the minimum and maximum FSC values. A histogram of the resolution values obtained from the conical FSC is shown in green. **e** Angular distribution of the resolution at FSC cutoff 0.5, generated using the 3DFSC file from Orientation Diagnostics job in CryoSPARC.

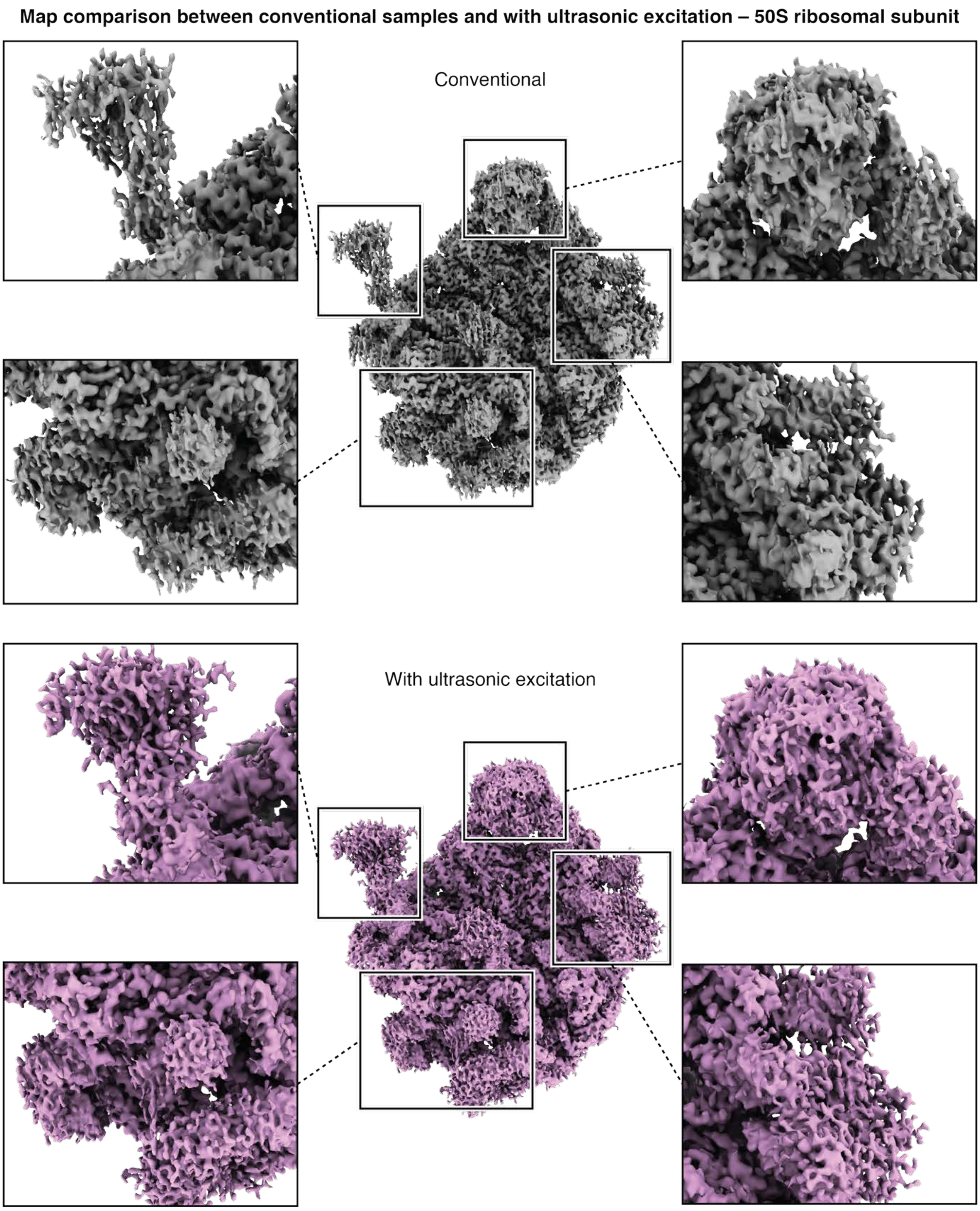


**Figure S4 | Comparison of the reconstructions from the conventional and ultrasonically excited samples of the 50S ribosomal subunit.** The streaky artefacts visible in the reconstruction from the conventional sample are caused by preferred orientation but are reduced with ultrasonic excitation. All maps are displayed at a level of 3σ above the mean.

**3 | Cryo-ET of a sample of the 50S ribosomal subunit**

Cryo-ET tilt series of a sample of the 50S ribosomal subunit (Fig. 2c) were collected at the Dubochet Center for Imaging in Lausanne, using a Thermo Fisher Titan Krios G4i transmission electron microscope equipped with a Falcon IV direct electron detector and SelectrisX energy filter. The data acquisition parameters are summarized in Table S1. The movies were motion-corrected by MotionCorr3[3] and imported into Aretomo2[4] for alignment and reconstruction. The resulting tomograms were imported into 3dmod[5] and Fiji2[6] to create the views presented in Fig. 2c.

**4 | Data processing and analysis – C-reactive protein**

High-resolution micrographs of a conventional cryo sample of human C-reactive protein as well as a sample that was subjected to ultrasonic excitation were collected at the Dubochet Center for Imaging in Lausanne, using a Thermo Fisher Titan Krios G4i transmission electron microscope equipped with a Falcon IV direct electron detector. Single-particle reconstructions were performed using CryoSPARC v4.7.1[2], with the corresponding workflows illustrated in Figs. S5-8. The data acquisition parameters and processing statistics are summarized in Table S3.

Micrographs were subjected to patch motion correction and patch CTF estimation. Micrographs with an estimated resolution of worse than 10 Å were rejected. Particles from the conventional and ultrasonically excited samples were picked using a blob picker with a diameter ranging from 20 to 130 Å. The particles were extracted with a box size of 384 px and Fourier-cropped to 96 px.

For the conventional dataset, particles were first subjected to six rounds of 2D classification during which particles corresponding to the C-reactive protein pentamer and decamer are separated. After 2D classification, particles corresponding to the conventional pentamer were subjected to *ab initio* reconstruction (one class, $C_1$ symmetry), homogeneous refinement ($C_5$ symmetry), and heterogeneous refinement (four classes, $C_5$ symmetry) using the previously homogeneously refined volume and three decoy volumes. Particles from the best class were then subjected to two rounds of 2D classification and re-extracted with a box size of 384 px. *Ab initio* reconstruction was then used on these remaining particles to produce three volumes ($C_5$ symmetry), each of which was then homogeneously refined ($C_5$ symmetry). Particles from the best two classes were subjected to a final round of heterogeneous refinement (four classes, $C_5$ symmetry). A random subset of 30,000 particles corresponding to the best volume in the prior heterogeneous refinement job were non-uniformly refined ($C_5$ symmetry) producing the final volume shown in Fig. S5.

Similarly, following 2D classification, particles corresponding to the conventional decamer were subjected to *ab initio* reconstruction (three classes, $C_1$ symmetry), homogeneous refinement ($C_1$ symmetry), and heterogeneous refinement (five classes, $C_1$ symmetry) using the previously homogeneously refined volume and four decoy volumes. Particles from the best class were then subjected to two rounds of 2D classification and re-extracted with a box size of 384 px. *Ab initio* reconstruction was then used on these remaining particles to produce three volumes ($C_1$ symmetry), each of which was then homogeneously refined ($C_1$ symmetry). Particles from the best two classes were subjected to a final round of heterogeneous refinement (four classes, $C_1$ symmetry). A random subset of 50,000 particles corresponding to the best volume in the prior heterogeneous refinement job were non-uniformly refined ($C_1$ symmetry) producing the final volume shown in Fig. S7.

The ultrasonically excited dataset was processed similarly. Particles were first subjected to six rounds of 2D classification after which particles corresponding to the ultrasonically excited pentamer were subjected to *ab initio* reconstruction (one class, $C_1$ symmetry), homogeneous refinement ($C_5$ symmetry), and heterogeneous refinement (five classes, $C_5$ symmetry) using the previously homogeneously refined volume and four decoy volumes. Particles from the best class were then subjected to two rounds of 2D classification and re-extracted with a box size of 384 px. *Ab initio* reconstruction was then used on these remaining particles to produce three volumes ($C_5$ symmetry), each of which was then homogeneously refined ($C_5$ symmetry). Particles from the best two classes were subjected to a final round of heterogeneous refinement (four classes, $C_5$ symmetry). A random subset of 30,000 particles corresponding to the best volume in the prior heterogeneous refinement job were non-uniformly refined ($C_5$ symmetry) producing the final volume shown in Fig. S6.

Following 2D classification, particles corresponding to the ultrasonically excited decamer were subjected to *ab initio* reconstruction (three classes, $C_1$ symmetry), homogeneous refinement ($C_1$ symmetry), and heterogeneous refinement (four classes, $C_1$ symmetry) using the previously homogeneously refined volume and three decoy volumes. Particles from the best class were then subjected to two rounds of 2D classification and re-extracted with a box size of 384 px. *Ab initio* reconstruction was then used on these remaining particles to produce three volumes ($C_1$ symmetry), each of which was then homogeneously refined ($C_1$ symmetry). Particles from the best two classes were subjected to a final round of heterogeneous refinement (four classes, $C_1$ symmetry). A random subset of 50,000 particles corresponding to the best volume in the prior heterogeneous refinement job were non-uniformly refined ($C_1$ symmetry) producing the final volume shown in Fig. S8.

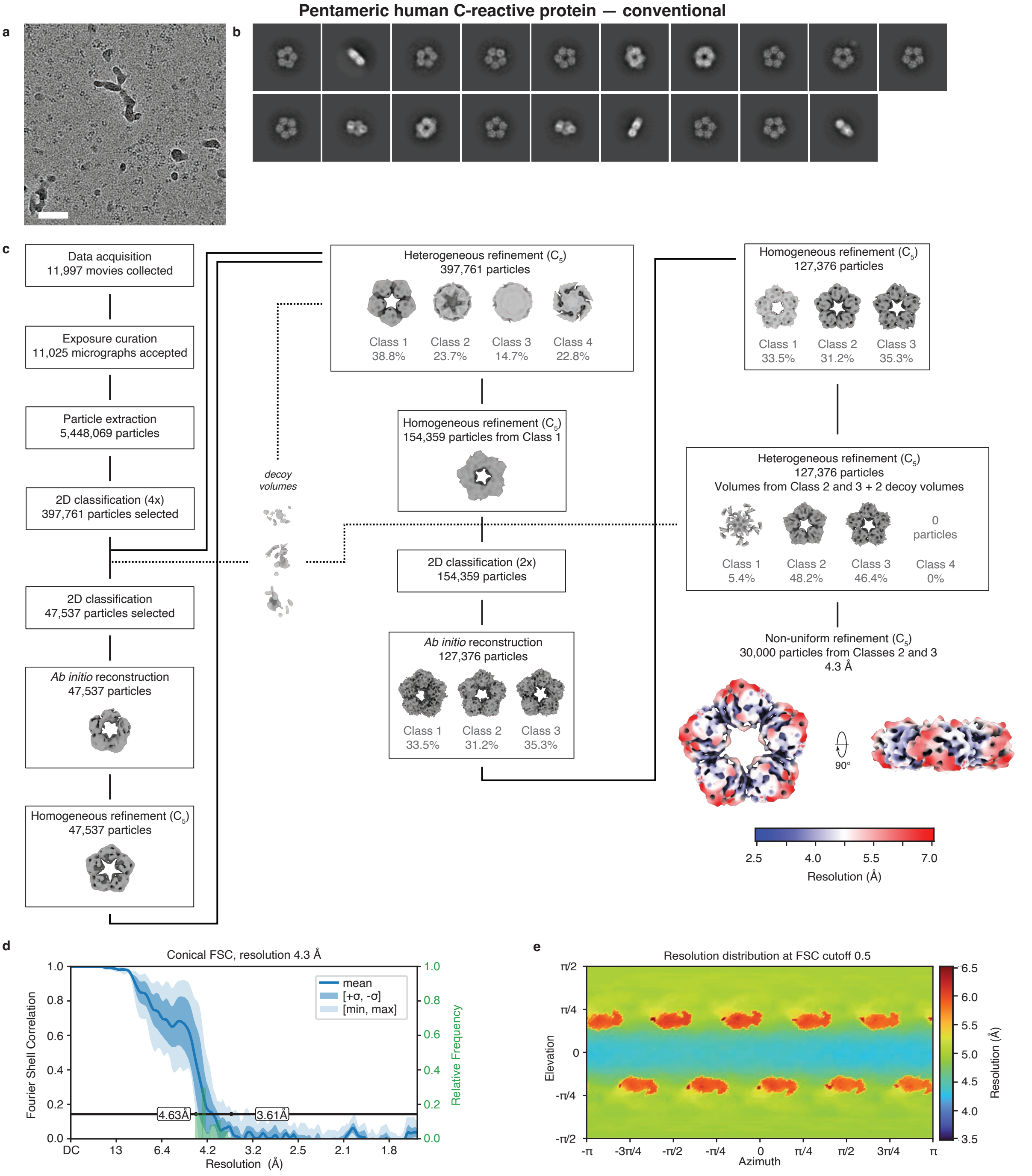


**Figure S5 | Processing workflow for the conventional sample of the C-reactive protein pentamer.** **a** Representative micrograph. Scale bar, 500 Å. **b** Selected 2D class averages. **c** Data processing workflow in CryoSPARC. The symmetry applied in each step is indicated in parentheses. The volumes are displayed at a level of 7σ above the mean. The final map is displayed at a level 10σ above the mean, with the local resolution estimation at the FSC cutoff of 0.5 indicated in color. **d** Conical FSC, with the 0.143 threshold indicated by a black line. The mean FSC value is shown as a solid blue line. Dark blue shading indicates one standard deviation of the FSC curve, while the light blue shading represents the minimum and maximum FSC values. A histogram of the resolution values obtained from the conical FSC is shown in green. **e** Angular distribution of the resolution at FSC cutoff 0.5, generated using the 3DFSC file from Orientation Diagnostics job in CryoSPARC.

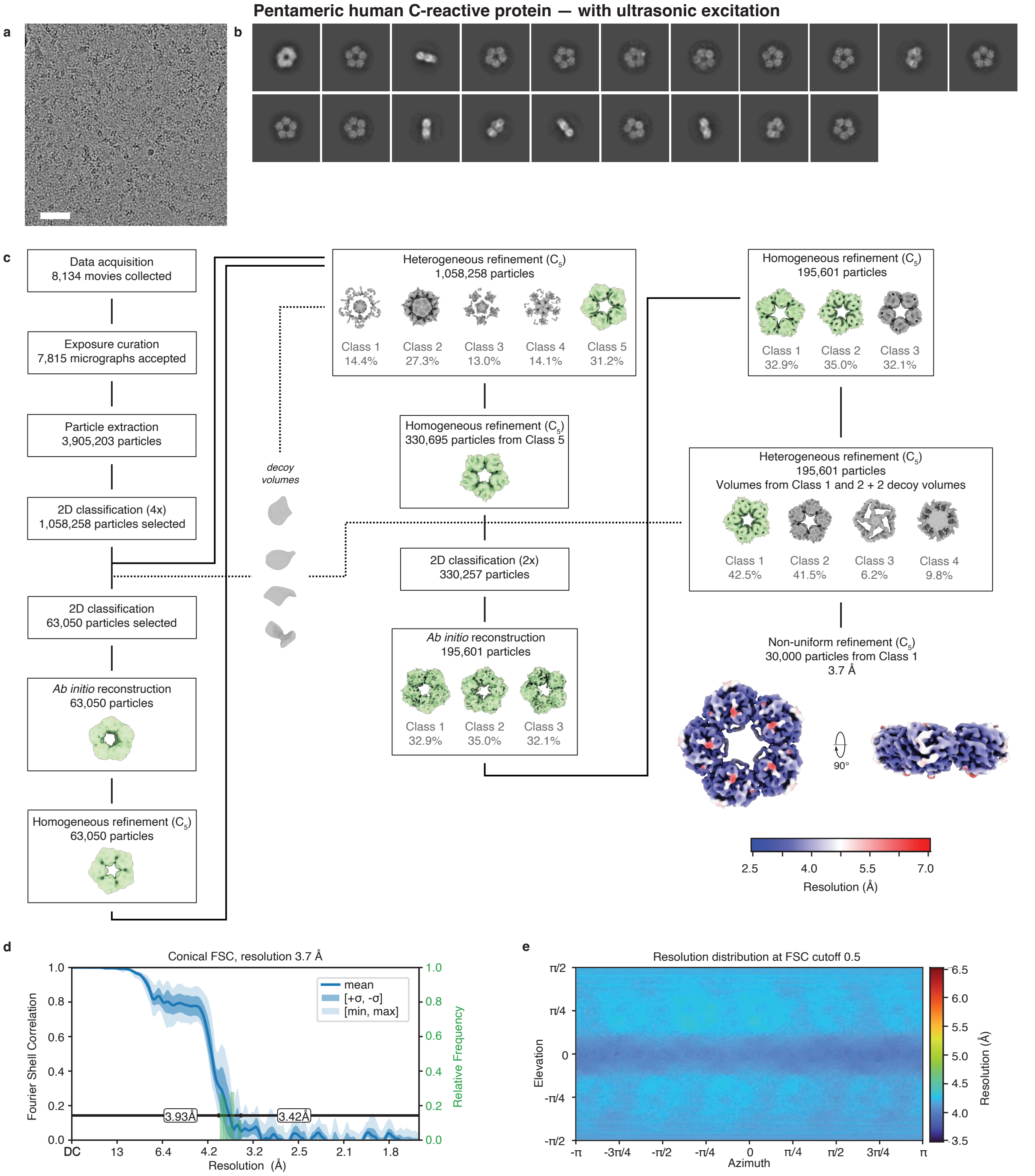


**Figure S6 | Processing workflow for the ultrasonically excited sample of the C-reactive protein pentamer.** **a** Representative micrograph. Scale bar, 500 Å. **b** Selected 2D class averages. **c** Data processing workflow in CryoSPARC. The symmetry applied in each step is indicated in parentheses. The volumes are displayed at a level of 7σ above the mean. The final map is displayed at a level 10σ above the mean, with the local resolution estimation at the FSC cutoff of 0.5 indicated in color. **d** Conical FSC, with the 0.143 threshold indicated by a black line. The mean FSC value is shown as a solid blue line. Dark blue shading indicates one standard deviation of the FSC curve, while the light blue shading represents the minimum and maximum FSC values. A histogram of the resolution values obtained from the conical FSC is shown in green. **e** Angular distribution of the resolution at FSC cutoff 0.5, generated using the 3DFSC file from Orientation Diagnostics job in CryoSPARC.

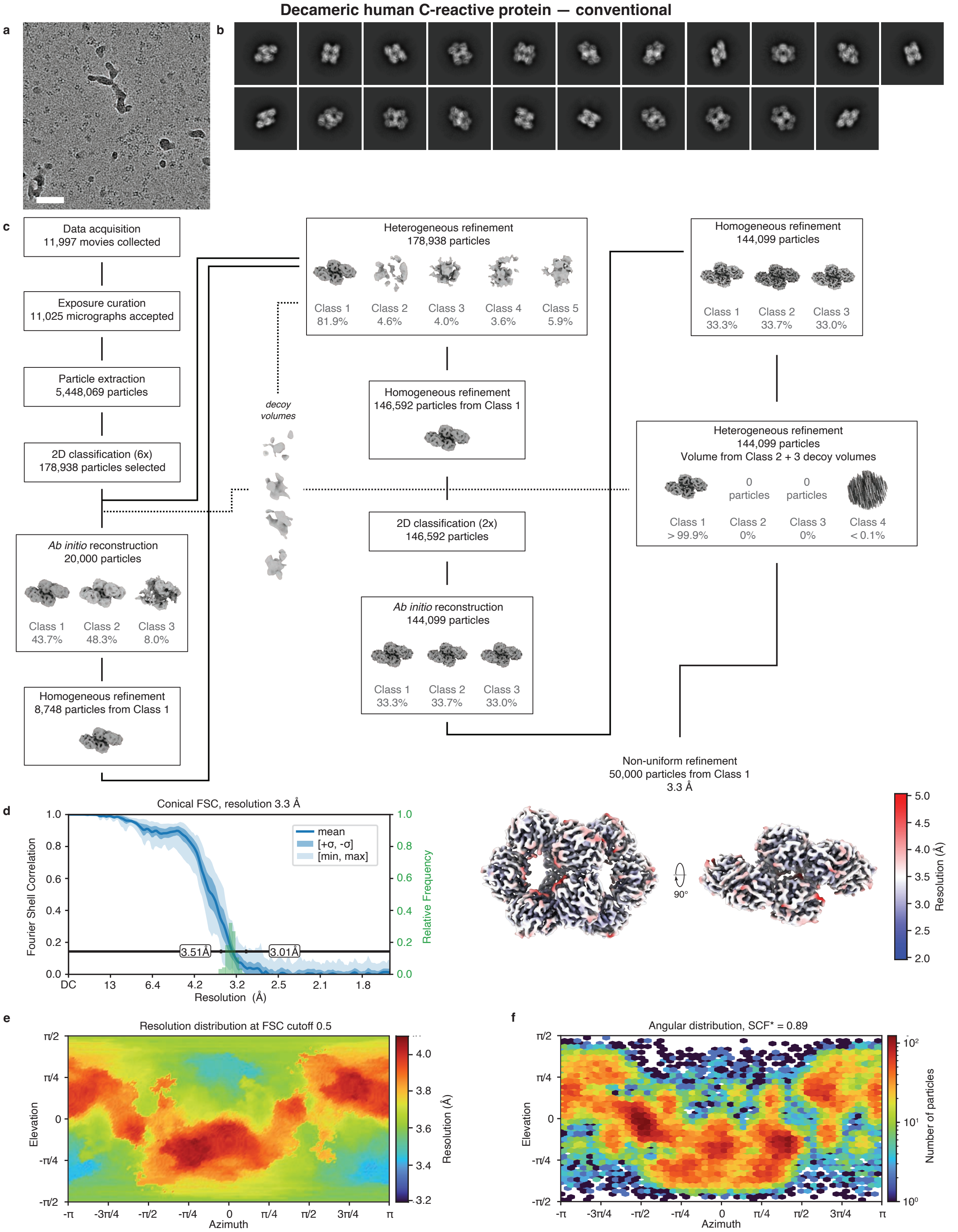


**Figure S7 | Processing workflow for the conventional sample of the C-reactive protein decamer.** **a** Representative micrograph. Scale bar, 500 Å. **b** Selected 2D class averages. **c** Data processing workflow in CryoSPARC. All reconstructions are performed with $C_1$ symmetry. The volumes are displayed at a level of 7σ above the mean. The final map is displayed at a level 10σ above the mean, with the local resolution estimation at the FSC cutoff of 0.5 indicated in color. **d** Conical FSC, with the 0.143 threshold indicated by a black line. The mean FSC value is shown as a solid blue line. Dark blue shading indicates one standard deviation of the FSC curve, while the light blue shading represents the minimum and maximum FSC values. A histogram of the resolution values obtained from the conical FSC is shown in green. **e** Angular distribution of the resolution at FSC cutoff 0.5, generated using the 3DFSC file from Orientation Diagnostics job in CryoSPARC. **f** Orientation distribution of the particles in the indicated final volume.

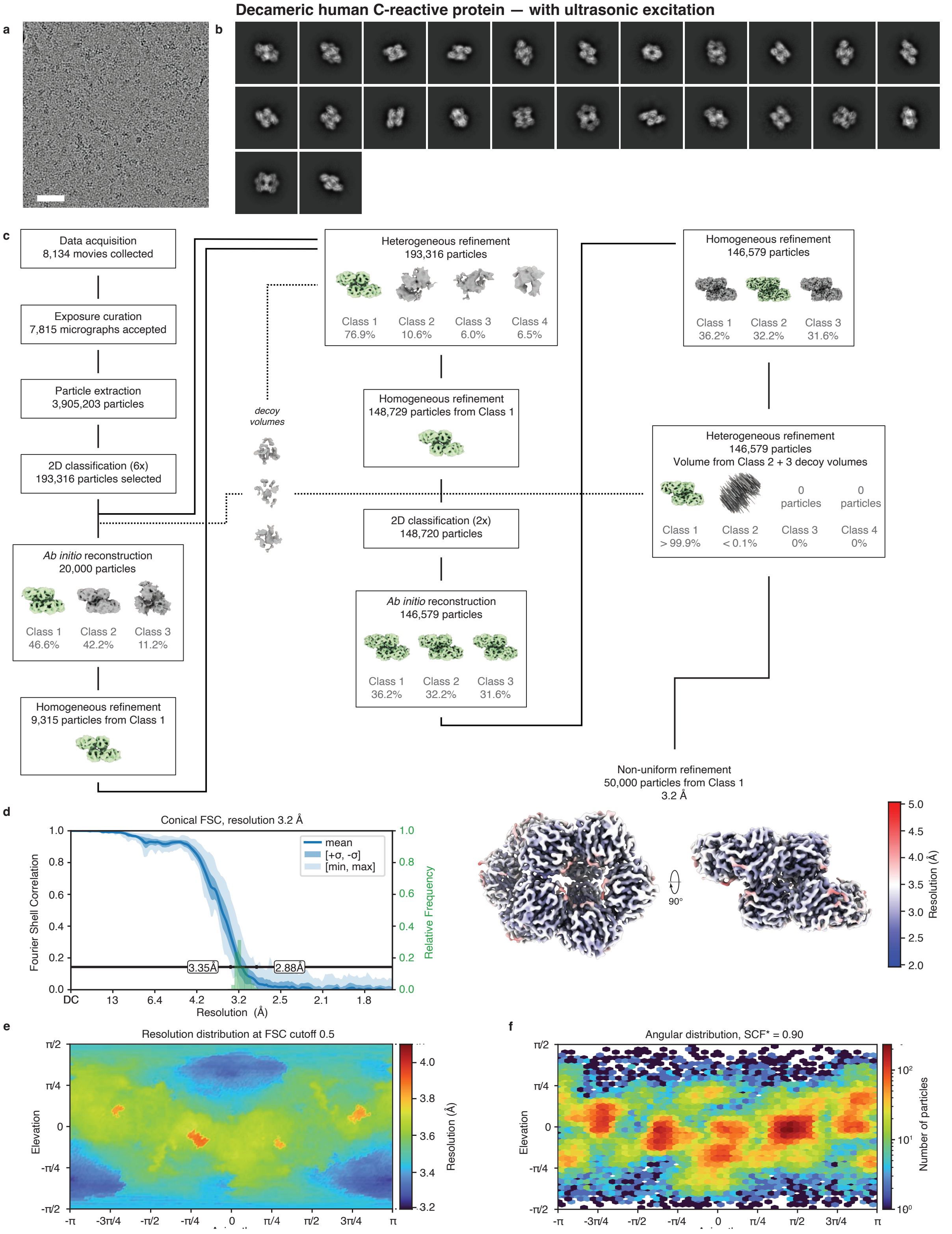


**Figure S8 | Processing workflow for the ultrasonically excited sample of the C-reactive protein decamer. a** Representative micrograph. Scale bar, 500 Å. **b** Selected 2D class averages. **c** Data processing workflow in CryoSPARC. All reconstructions are performed with $C_1$ symmetry. The volumes are displayed at a level of 7σ above the mean. The final map is displayed at a level 10σ above the mean, with the local resolution estimation at the FSC cutoff of 0.5 indicated in color. **d** Conical FSC, with the 0.143 threshold indicated by a black line. The mean FSC value is shown as a solid blue line. Dark blue shading indicates one standard deviation of the FSC curve, while the light blue shading represents the minimum and maximum FSC values. A histogram of the resolution values obtained from the conical FSC is shown in green. **e** Angular distribution of the resolution at FSC cutoff 0.5, generated using the 3DFSC file from Orientation Diagnostics job in CryoSPARC. **f** Orientation distribution of the particles in the indicated final volume.

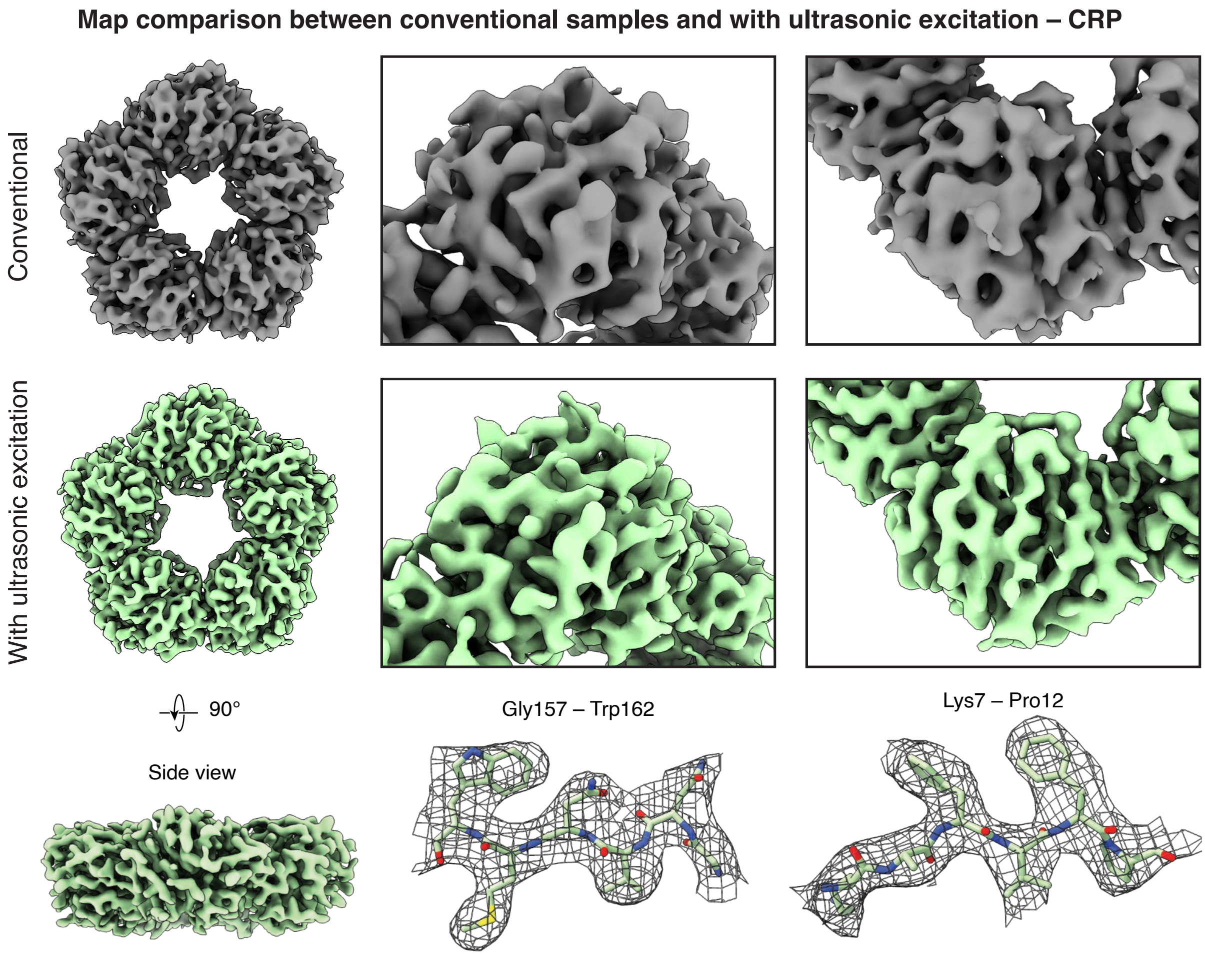


**Figure S9 | Map comparison between the conventional and ultrasonically excited maps for the pentameric C-reactive protein.** To illustrate artifacts in the map obtained from the conventional sample of the pentameric C-reactive protein, top views (left) for the conventional (top, in grey) and ultrasonically excited (bottom, in green) samples are shown. Details of these maps reveal artifacts in the conventional map as a result of preferred orientation. Further, to demonstrate high-resolution and interpretability of the cryo-EM map shown in the bottom panel, a model (PDB ID: 7PKD[7]) is docked and jiggle-fitted within the map. Several sections of the amino acid backbone are shown (right). All maps are displayed with a level at 7.5$\sigma$ above the mean.

**5 | Data processing and analysis – hemagglutinin**

High-resolution micrographs of a conventional cryo sample of hemagglutinin as well as a sample that was subjected to ultrasonic excitation were collected at the Dubochet Center for Imaging in Lausanne, using a Thermo Fisher Titan Krios G4i transmission electron microscope equipped with a Falcon IV direct electron detector. Single-particle reconstructions were performed using CryoSPARC v4.7.1[2], with the corresponding workflows illustrated in Figs. S10 and S11. The data acquisition parameters and processing statistics are summarized in Table S4.

Micrographs were subjected to patch motion correction and patch CTF estimation. Micrographs with an estimated resolution of worse than 10 Å were rejected. Particles from the conventional and ultrasonically excited samples were picked using a blob picker with a diameter ranging from either 50 to 150 Å (conventional sample) or 80 to 150 Å (ultrasonically excited sample). The particles were extracted with a box size of 384 px and initially Fourier-cropped to 96 px.

For the conventional dataset, particles were first subjected to four rounds of 2D classification followed by *ab initio* reconstruction (three classes, $C_1$ symmetry) and then two rounds of heterogeneous refinement (all three *ab initio* classes, $C_3$ symmetry). Particles from the best class resulting from the final round of heterogeneous refinement were re-extracted with a box size of 384 px and non-uniformly refined ($C_3$ symmetry) producing the final volume shown in Fig. S10.

Particles from the ultrasonically excited dataset were first subjected to three rounds of 2D classification. A comparison of the obtained 2D classes is shown in Fig. S12 highlighting that ultrasonic excitation populates side views that are absent in the conventional dataset. Following 2D classification, top/bottom views and side views of the hemagglutinin trimer were separated. *Ab initio* reconstruction ($C_1$ symmetry) was first performed on a subset of particles containing predominantly top/bottom views producing a truncated hemagglutinin trimer. Separately, *ab initio* reconstruction ($C_1$ symmetry) was performed on particles representing predominantly side views producing a volume closely resembling the hemagglutinin trimer. Particles from this second *ab initio* reconstruction were then subjected to homogeneous refinement ($C_3$ symmetry), followed by two rounds of heterogeneous refinement (three classes, $C_3$ symmetry) using the previously homogeneously refined volume and two decoys. Particles from the best class from the final round of heterogeneous refinement were then subjected to homogeneous refinement ($C_3$ symmetry) and subsequently re-extracted with a box size of 384 px. All originally selected particles were then heterogeneously refined ($C_3$ symmetry) against five copies of the *ab initio* reconstruction derived from the top/bottom views and one copy of a hemagglutinin volume derived from the side views. Particles from the best class were then homogeneously refined ($C_3$ symmetry) and subjected to a further round of heterogeneous refinement ($C_3$ symmetry) using three volumes: one copy of the *ab initio* reconstruction derived from the top/bottom views, and two copies of the complete hemagglutinin volume generated in the previous round of homogeneous refinement. Particles from the best class resulting from the final round of heterogeneous refinement were non-uniformly refined ($C_3$ symmetry) producing the final volume shown in Fig. 3b and Fig. S11. Here, best class refers to the reconstruction with no indicators of directional anisotropy (e.g. missing and density). Nevertheless, to enable a comparison of angular distributions after data processing, the other two classes in the final round of heterogeneous refinement were also re-extracted and non-uniformly refined and are displayed in Fig. S11.

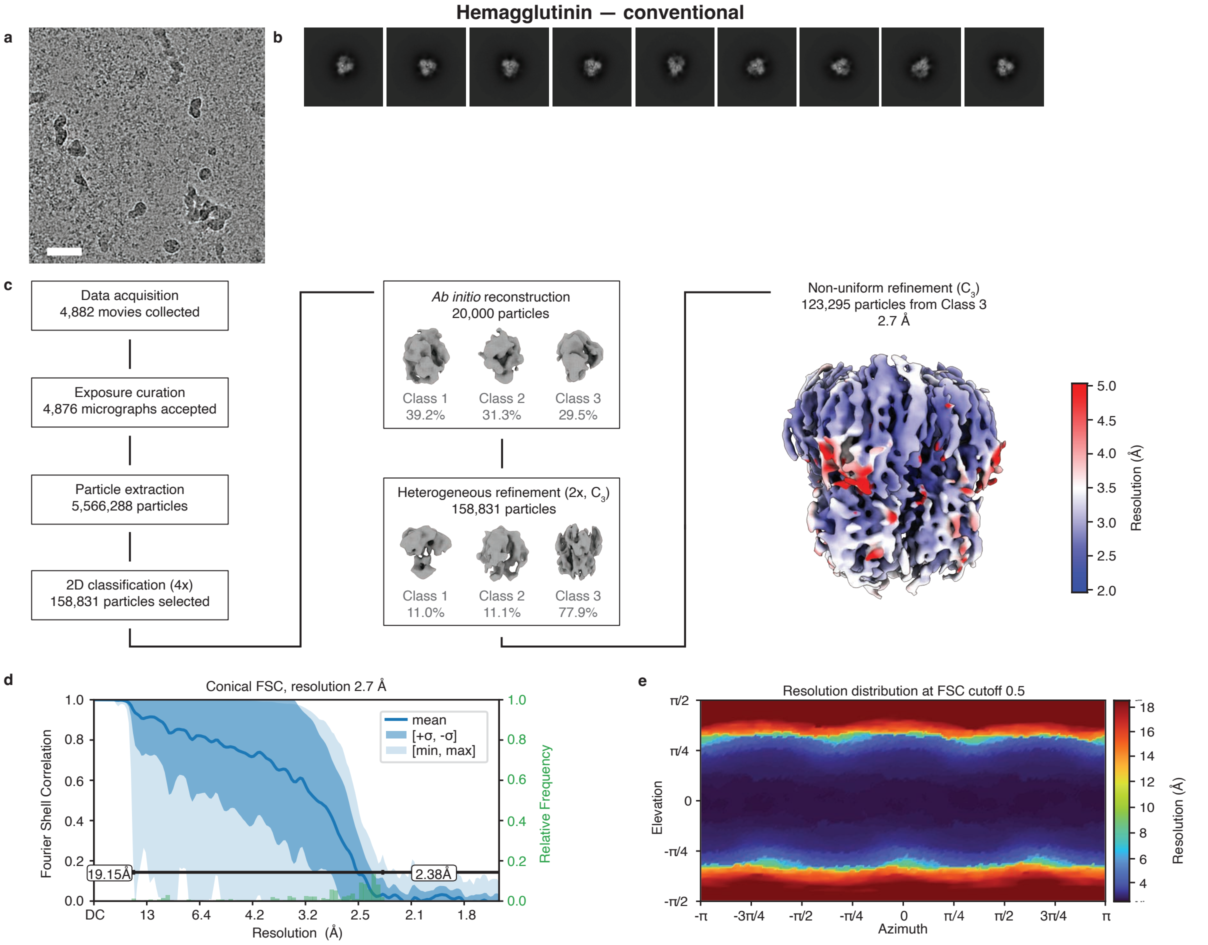


**Figure S10 | Cryo-EM processing workflow for the conventional sample of hemagglutinin.** **a** Representative micrograph. Scale bar, 500 Å. **b** Selected 2D class averages. **c** Data processing workflow in CryoSPARC. Symmetry imposed is indicated in parentheses. All volumes are displayed at a level of 10σ above the mean. The final map is colored according to the estimated local resolution at the FSC cutoff of 0.5. **d** Conical FSC, with the 0.143 threshold indicated by a black line. The mean FSC value is shown as a solid blue line. Dark blue shading indicates one standard deviation of the FSC curve, while the light blue shading represents the minimum and maximum FSC values. A histogram of the resolution values obtained from the conical FSC is shown in green. **e** Angular distribution of the resolution at FSC cutoff 0.5, generated using the 3DFSC file from Orientation Diagnostics job in CryoSPARC.

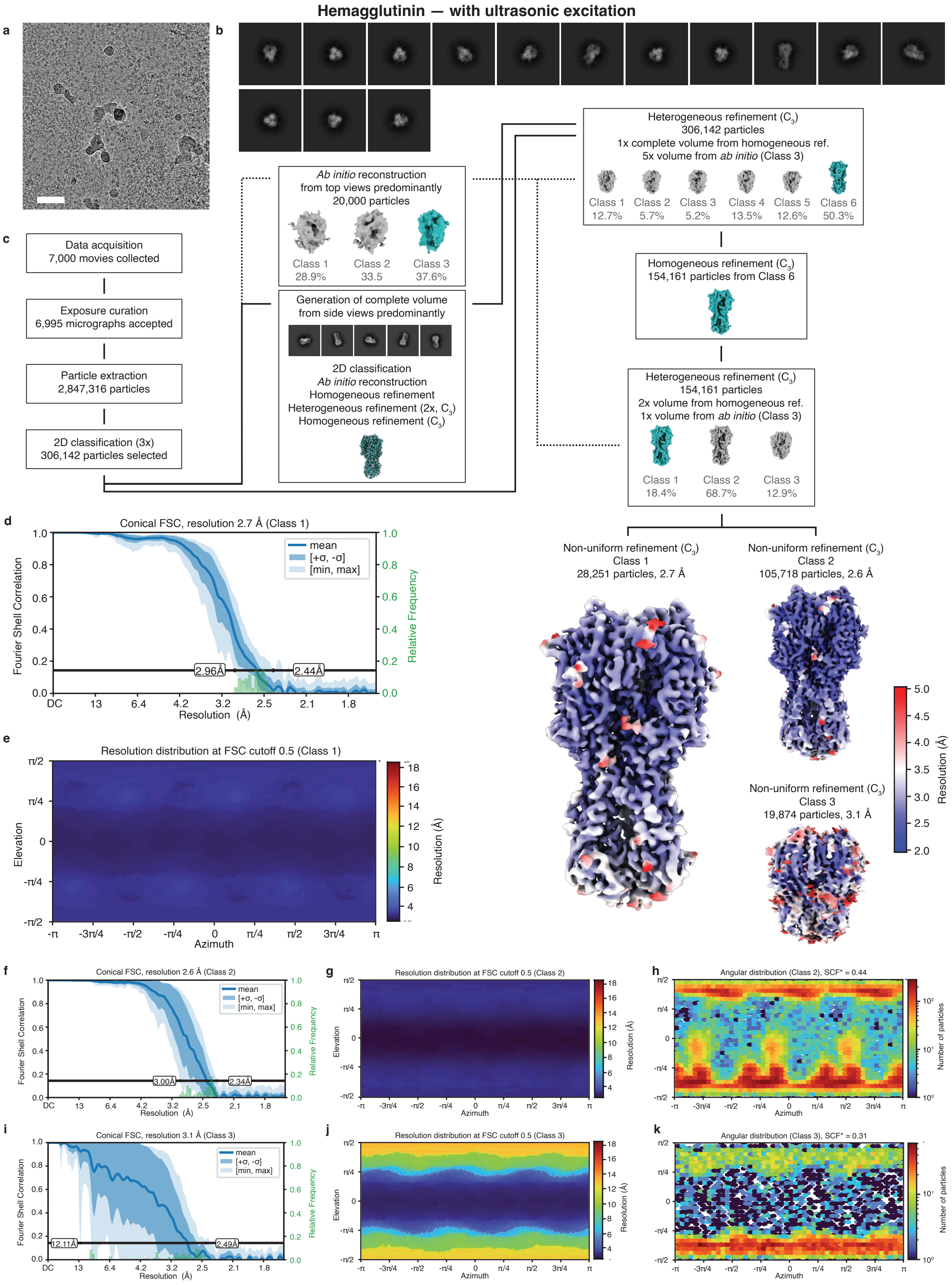
Hemagglutinin — with ultrasonic excitation
a
b
c
Data acquisition
7,000 movies collected
Exposure curation
6,995 micrographs accepted
Particle extraction
2,847,316 particles
2D classification (3x)
306,142 particles selected
Ab initio reconstruction
from top views predominantly
20,000 particles
Class 1 28.9%
Class 2 33.5
Class 3 37.6%
Generation of complete volume
from side views predominantly
2D classification
Ab initio reconstruction
Homogeneous refinement
Heterogeneous refinement (2x, C3)
Homogeneous refinement (C3)
Heterogeneous refinement (C3)
306,142 particles
1x complete volume from homogeneous ref.
5x volume from ab initio (Class 3)
Class 1 12.7%
Class 2 5.7%
Class 3 5.2%
Class 4 13.5%
Class 5 12.6%
Class 6 50.3%
Homogeneous refinement (C3)
154,161 particles from Class 6
Heterogeneous refinement (C3)
154,161 particles
2x volume from homogeneous ref.
1x volume from ab initio (Class 3)
Class 1 18.4%
Class 2 68.7%
Class 3 12.9%
Non-uniform refinement (C3)
Class 1
28,251 particles, 2.7 Å
Non-uniform refinement (C3)
Class 2
105,718 particles, 2.6 Å
Non-uniform refinement (C3)
Class 3
19,874 particles, 3.1 Å
Resolution (Å)
d
Conical FSC, resolution 2.7 Å (Class 1)
Fourier Shell Correlation
Relative Frequency
Resolution (Å)
mean
[+σ, -σ]
[min, max]
2.96Å
2.44Å
e
Resolution distribution at FSC cutoff 0.5 (Class 1)
Elevation
Azimuth
Resolution (Å)
f
Conical FSC, resolution 2.6 Å (Class 2)
3.00Å
2.34Å
g
Resolution distribution at FSC cutoff 0.5 (Class 2)
h
Angular distribution (Class 2), SCF* = 0.44
Number of particles
i
Conical FSC, resolution 3.1 Å (Class 3)
12.11Å
2.49Å
j
Resolution distribution at FSC cutoff 0.5 (Class 3)
k
Angular distribution (Class 3), SCF* = 0.31

**Figure S11 | Processing workflow for the ultrasonically excited sample of hemagglutinin.** **a** Representative micrograph. Scale bar, 500 Å. **b** Selected 2D class averages. **c** Data processing workflow in CryoSPARC. Symmetry imposed is indicated in parentheses. All volumes are displayed at a level of 10$\sigma$ above the mean. Final maps are colored according to the estimated local resolution at the FSC cutoff of 0.5. **d,f,i** Conical FSC, with the 0.143 threshold indicated by a black line. The mean FSC value is shown as a solid blue line. Dark blue shading indicates one standard deviation of the FSC curve, while the light blue shading represents the minimum and maximum FSC values. A histogram of the resolution values obtained from the conical FSC is shown in green. **e,g,j** Angular distribution of the resolution at FSC cutoff 0.5, generated using the 3DFSC file from Orientation Diagnostics job in CryoSPARC. **h,k** Orientation distribution of the particles in the indicated final volume.

**Ultrasonic excitation during vitrification populates side views of hemagglutinin**

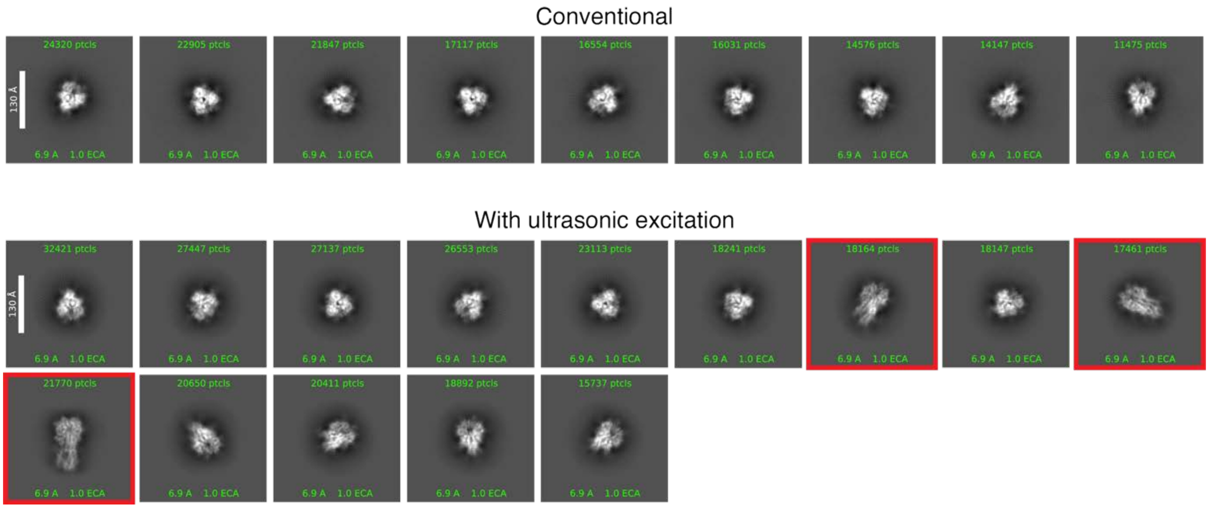


**Figure S12 | Ultrasonic excitation during vitrification populates side views of hemagglutinin.** With ultrasonic excitation, the 2D class averages show side views (bottom, highlighted in red), which are missing in a conventional sample (top).

**6 | Estimate of the angular distribution in the conventional and ultrasonically excited hemagglutinin samples**

Figure 3 compares reconstructions of hemagglutinin from a conventional sample as well as a sample that was subjected to ultrasonic excitation during vitrification, with the corresponding workflows for obtaining the reconstructions detailed in Supplementary Information 5. The angular distributions displayed in Fig. 3 represent the distributions of the particles that contribute to the maps displayed. However, because hemagglutinin has such strong preferred orientation, these distributions likely do not accurately reflect the angular distribution of all particles in the sample. The conventional dataset only yielded a truncated reconstruction, so that the particle orientations determined against this map likely have large errors. For the sample vitrified with ultrasonic excitation, the workflow that yields the volume shown in the main text removes a large number of top and bottom views during the heterogeneous refinement steps, so their population is likely underrepresented in the angular distribution.

To arrive at a more realistic estimate of the angular distribution, we therefore refined all particles picked after 2D classification against the complete volume of hemagglutinin shown in Fig. 3b. The corresponding workflows are illustrated in Fig. S13a. For the conventional sample, this produces a volume that more closely resembles the hemagglutinin trimer but still exhibits a large amount of directional anisotropy. The corresponding angular distribution is largely dominated by top and bottom views and has an SCF* value of 0.43 (Fig. S13b). For the sample vitrified with ultrasonic excitation, the same process yields an angular distribution that additionally features a significant population of side views and has an SCF* of 0.56. Note, this procedure for estimating the angular distribution of the sample likely includes a considerable fraction of damaged particles in the statistics that would otherwise be removed through heterogeneous refinement. Such particles likely have a large error in the angular assignment, which has the effect of artificially increasing the SCF* value.

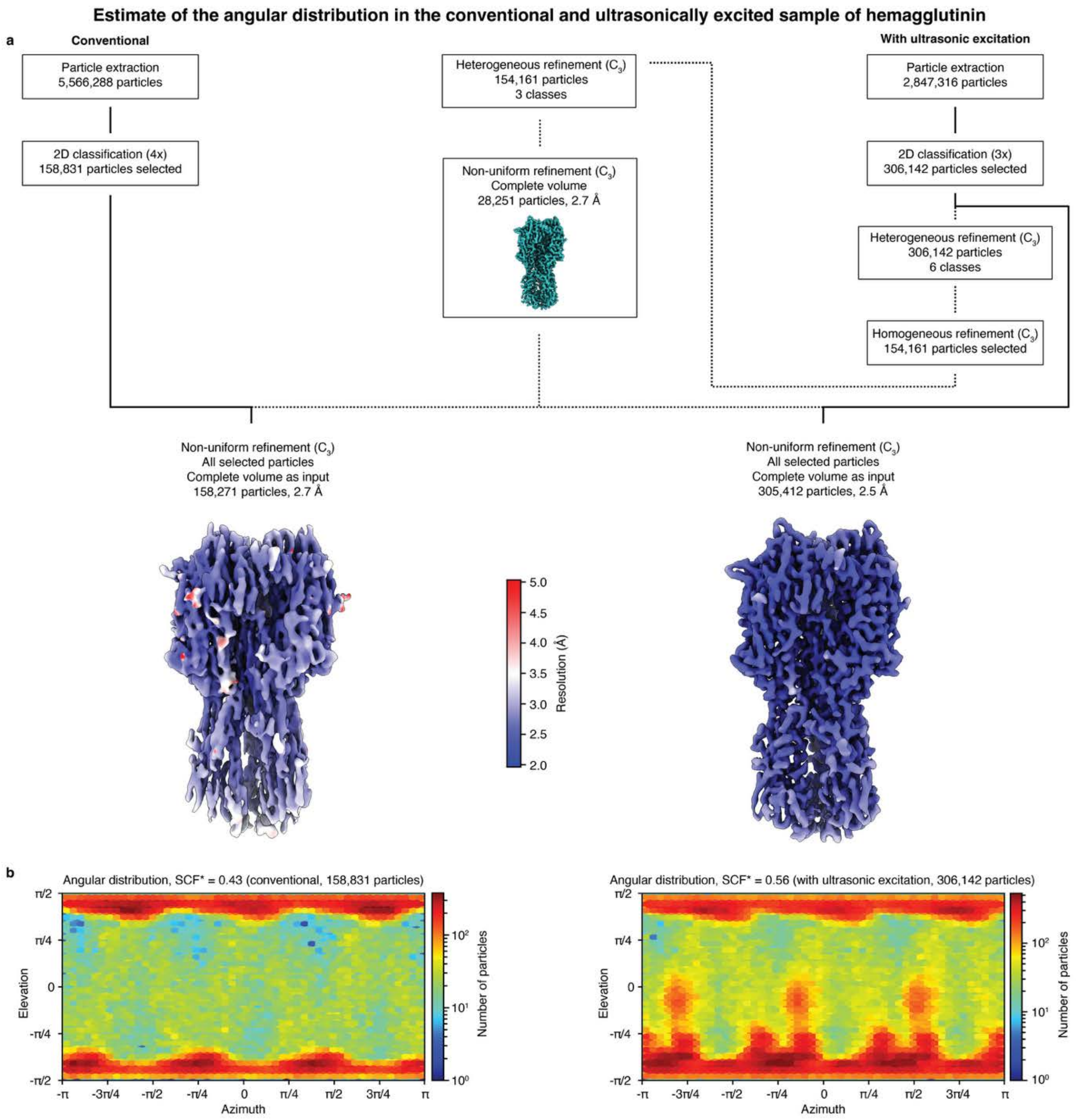


**Figure S13 | Estimate of the angular distribution in the conventional and ultrasonically excited sample of hemagglutinin. a** Data processing workflow in CryoSPARC. All maps are displayed at the level of 10σ above the mean. Symmetry imposed is indicated in parentheses. The final map is displayed with the local resolution estimation indicated in color. **b** Estimate of the angular distribution obtained from a non-uniform refinement of all selected particles against the complete hemagglutinin volume from Fig. 3b.

### 7 | Proof-of-concept implementation of ultrasonic excitation in a plunge-freezing device

We demonstrate a proof-of-principle implementation of ultrasonic excitation in a plunge-freezing device as illustrated in Fig. S14a. Samples of viral hemagglutinin (unclipped grids) are plunge-frozen in liquid ethane with our modified Thermo Fisher Vitrobot Mark IV (Supplementary Information 1). Four ultrasonic transducers (SECO SC049) are placed below the climate chamber to excite the sample with ultrasonic waves before it plunges into the cryogen. The transducers, which are driven with a 490 kHz square wave of 150 V amplitude, point towards the surface of the cryogen and are located at a distance of 9 mm from the path along which the sample travels.

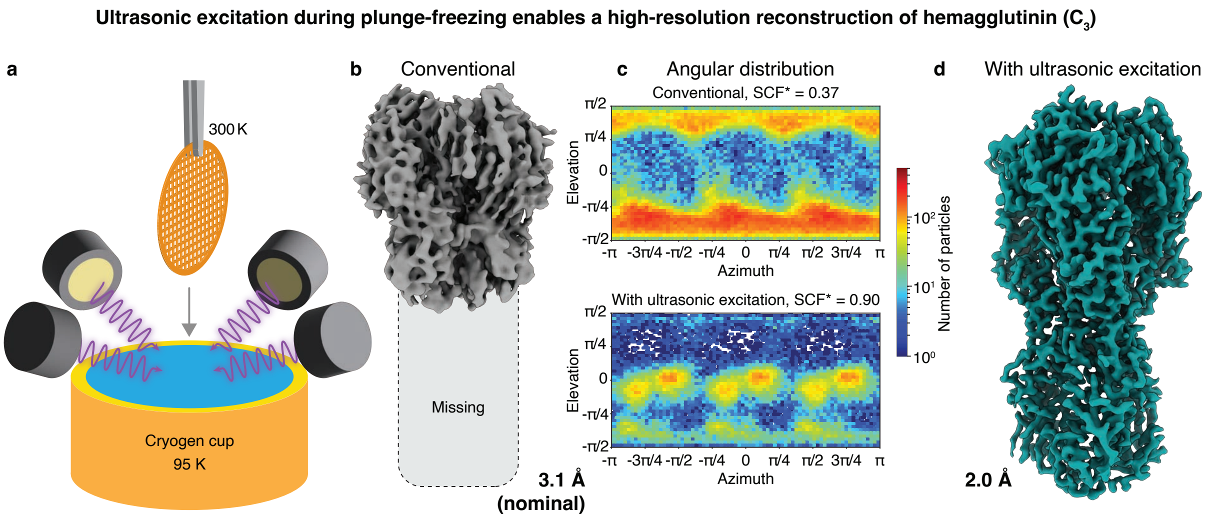


**Figure S14 | Proof-of-concept implementation of ultrasonic excitation in a plunge-freezing device. a** Experimental concept. A cryo-EM sample is prepared by plunge freezing while mechanical oscillations of the specimen support are excited with four ultrasonic transducers. **b–d** Reconstructions and angular distributions from a conventional, plunge-frozen hemagglutinin sample as well as a sample that was excited with ultrasonic waves during plunge-freezing. Because of strong preferred orientation, the conventional sample yields an only a truncated reconstruction (**b,c**). In contrast, ultrasonic excitation improves the angular distribution by populating side views, allowing us to obtain a complete volume at 2.0 Å resolution (**c,d**).

High-resolution micrographs of a conventional cryo sample of hemagglutinin as well as a sample that was subjected to ultrasonic excitation during plunging were collected at the Dubochet Center for Imaging in Lausanne, using a Thermo Fisher Titan Krios G4i transmission electron microscope equipped with a Falcon IV direct electron detector. Single-particle reconstructions were performed using CryoSPARC v4.7.1[2], with the corresponding workflows illustrated in Figs. S15 and S16. The data acquisition parameters and processing statistics are summarized in Table S5.

Micrographs were subjected to patch motion correction and patch CTF estimation. Micrographs with an estimated resolution of worse than 10 Å were rejected. Particles from the conventional and ultrasonically excited samples were picked using a blob picker with a diameter ranging from 80 to 150 Å. The particles were extracted with a box size of 384 px and initially Fourier-cropped to 96 px.

For the conventional dataset, particles were first subjected to three rounds of 2D classification followed by *ab initio* reconstruction (three classes, $C_1$ symmetry) and then two rounds of heterogeneous refinement (the highest quality *ab initio* class and three decoys, $C_3$ symmetry). Particles from the best class were homogeneously refined ($C_3$ symmetry), the resulting volume and particles were then subjected to one further round of heterogeneous refinement with three decoy volumes ($C_3$ symmetry). Particles from the best class resulting from the final round of heterogeneous refinement were re-extracted with a box size of 384 px and non-uniformly refined ($C_3$ symmetry) producing the final volume shown in Fig. S14b and S15.

Particles from the ultrasonically excited dataset were first subjected to one round of 2D classification. Following 2D classification, top/bottom views and side views of the hemagglutinin trimer were separated. *Ab initio* reconstruction ($C_1$ symmetry) was first performed on a subset of particles containing predominantly side views producing a volume closely resembling the hemagglutinin trimer. Good particles were then subjected to two rounds of heterogeneous refinement using two good volumes and two decoy volumes ($C_3$ symmetry). Particles from the best class from the second round of heterogeneous refinement were then non-uniformly refined ($C_3$ symmetry) and subjected to a final round of heterogeneous refinement using three copies of the volume from the non-uniform refinement job were used ($C_3$ symmetry). Particles from the best class resulting from the final round of heterogeneous refinement were non-uniformly refined ($C_3$ symmetry) producing the final volume shown in Fig. S14d and Fig. S16. Here, best class refers to the reconstruction with the highest resolution and no indicators of directional anisotropy. Nevertheless, to enable a comparison of angular distributions after data processing, the other two classes in the final round of heterogeneous refinement were also re-extracted with a box size of 384 px and non-uniformly refined ($C_3$ symmetry) and are displayed in Fig. S16.

We follow the procedure detailed in Supplementary Information 6 to arrive at realistic estimates of the angular distributions of the conventional, plunge-frozen sample as well as the sample plunge-frozen with ultrasonic excitation. We refine all particles picked after 2D classification against the complete volume of hemagglutinin shown in Fig. S14d. The corresponding workflows are illustrated in Fig. S17a. For the conventional, plunge-frozen sample, this produces a volume that more closely resembles the hemagglutinin trimer but still exhibits a large amount of directional anisotropy. The corresponding angular distribution is largely dominated by top and bottom views and has an SCF* value of 0.46 (Fig. S17b). For the sample plunge-frozen with ultrasonic excitation, the same process yields an angular distribution that additionally features a significant population of side views and has an SCF* value of 0.74.

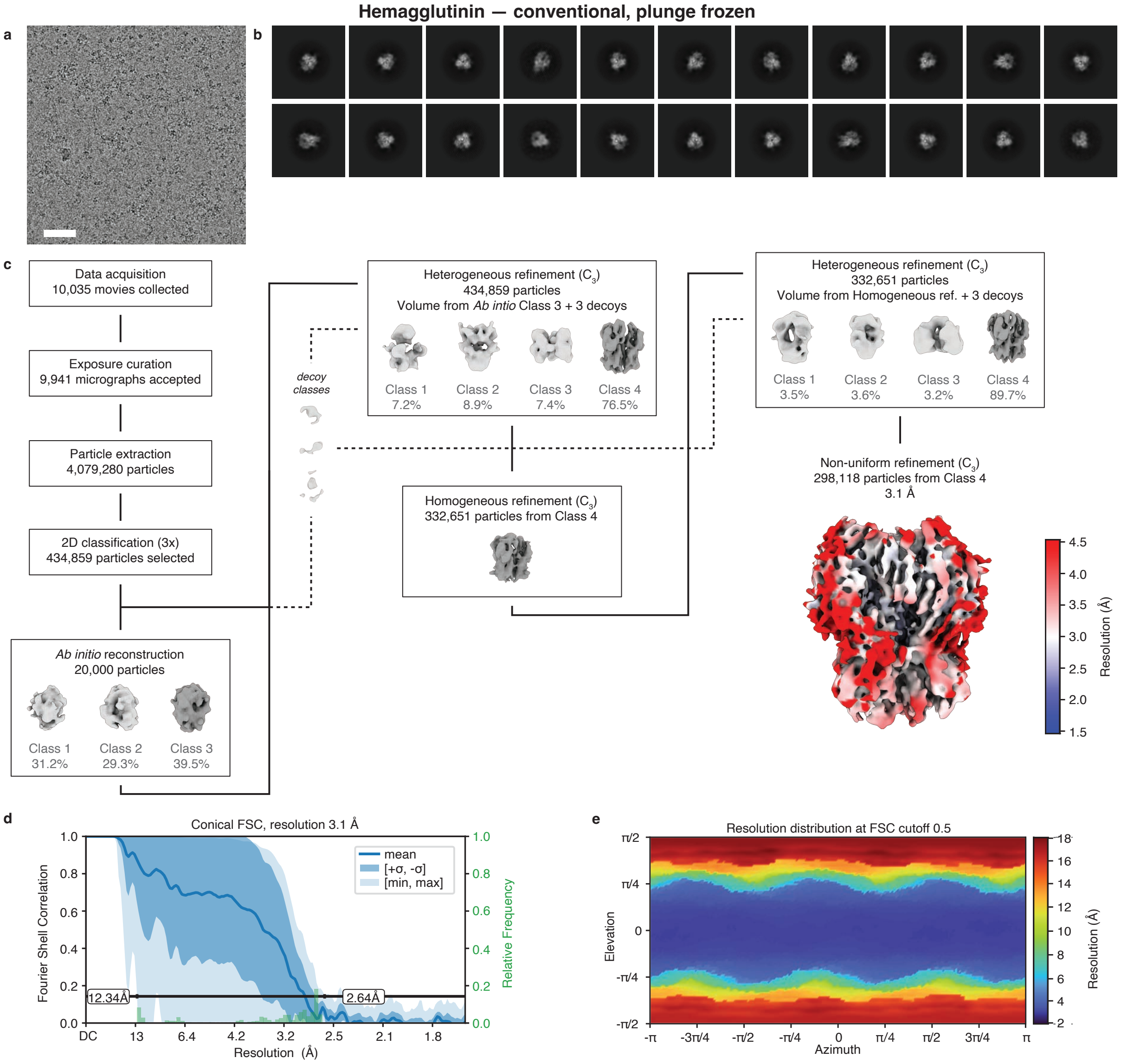


**Figure S15 | Cryo-EM processing workflow for the conventional, plunge-frozen sample of hemagglutinin.** **a** Representative micrograph. Scale bar, 500 Å. **b** Selected 2D class averages. **c** Data processing workflow in CryoSPARC. Symmetry imposed is indicated in parentheses. All volumes are displayed at a level of 10σ above the mean. The final map is colored according to the estimated local resolution at the FSC cutoff of 0.5. **d** Conical FSC, with the 0.143 threshold indicated by a black line. The mean FSC value is shown as a solid blue line. Dark blue shading indicates one standard deviation of the FSC curve, while the light blue shading represents the minimum and maximum FSC values. A histogram of the resolution values obtained from the conical FSC is shown in green. **e** Angular distribution of the resolution at FSC cutoff 0.5, generated using the 3DFSC file from Orientation Diagnostics job in CryoSPARC.

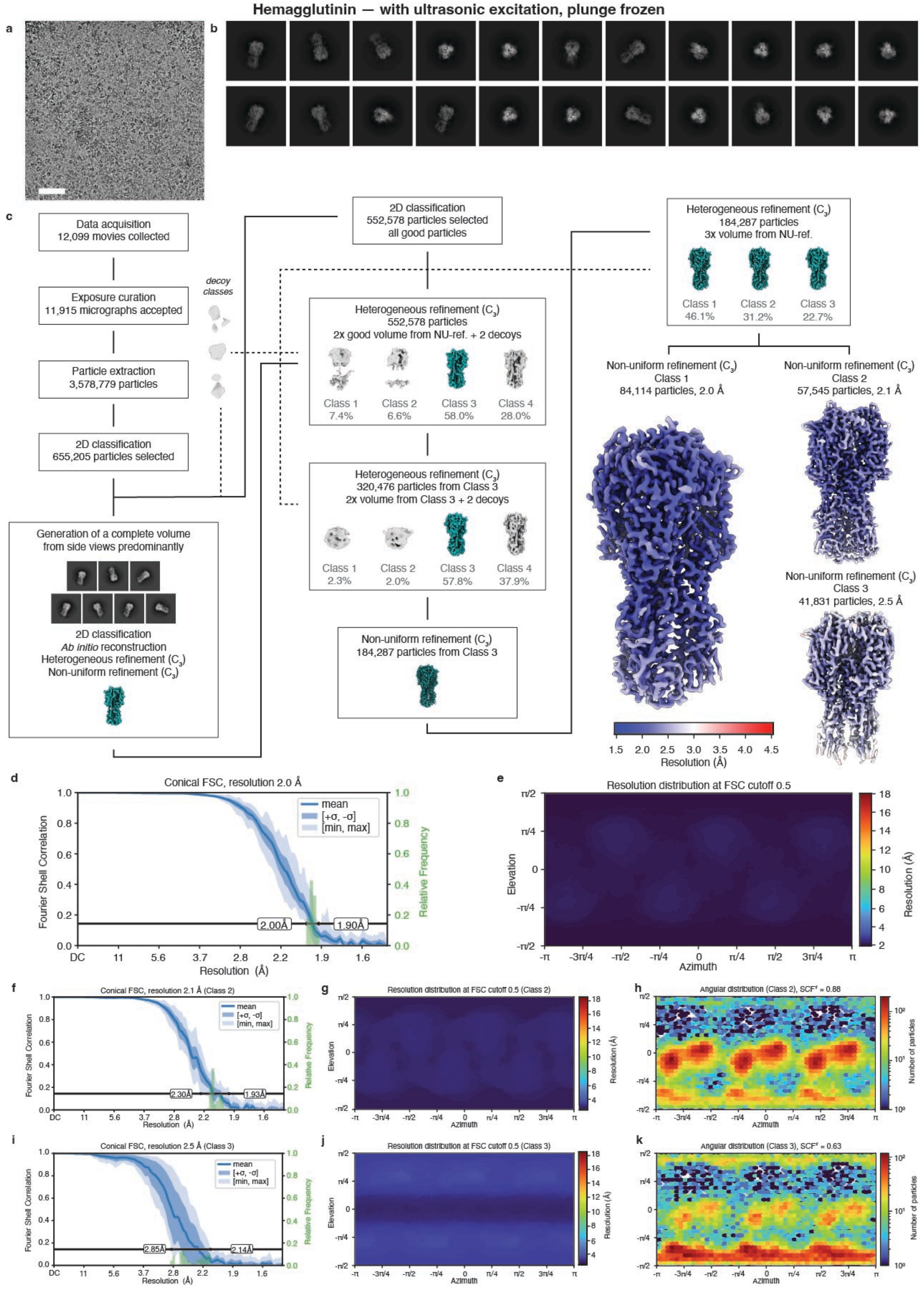


**Figure S16 | Processing workflow for the ultrasonically excited, plunge-frozen sample of hemagglutinin. A** Representative micrograph. Scale bar, 500 Å. **B** Selected 2D class averages. **C** Data processing workflow in CryoSPARC. Symmetry imposed is indicated in parentheses. All volumes are displayed at a level of 10$\sigma$ above the mean. Final maps are colored according to the estimated local resolution at the FSC cutoff of 0.5. **d,f,i** Conical FSC, with the 0.143 threshold indicated by a black line. The mean FSC value is shown as a solid blue line. Dark blue shading indicates one standard deviation of the FSC curve, while the light blue shading represents the minimum and maximum FSC values. A histogram of the resolution values obtained from the conical FSC is shown in green. **e,g,j** Angular distribution of the resolution at FSC cutoff 0.5, generated using the 3DFSC file from Orientation Diagnostics job in CryoSPARC. **h,k** Orientation distribution of the particles in the indicated final volume.

### Estimate of the angular distribution in the conventional and ultrasonically excited sample of hemagglutinin, plunge frozen

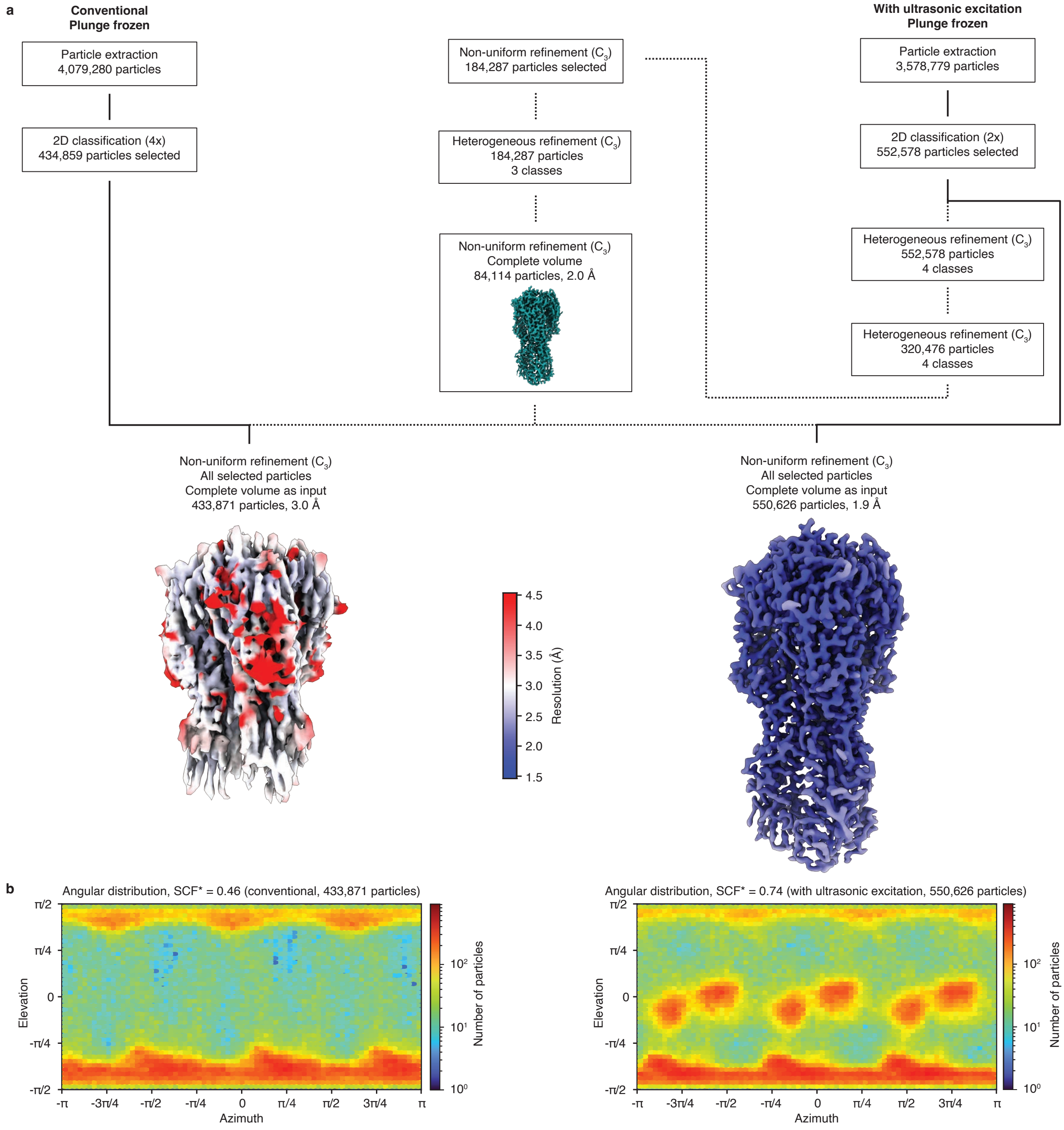


**Figure S17 | Estimate of the angular distribution in the conventionally plunge-frozen hemagglutinin sample the sample plunge-frozen with ultrasonic excitation. a** Data processing workflow in CryoSPARC. All maps are displayed at a level of 10σ above the mean. Symmetry imposed is indicated in parentheses. The final map is displayed with the local resolution estimation indicated in color. **b** Estimate of the angular distribution obtained from a non-uniform refinement of all selected particles against the complete hemagglutinin volume from Fig. S13a,d.

**8 | Cryo-ET data collection statistics**

**Supplementary Table 1:** Data collection parameters for the 50S ribosomal subunit tilt series.

| | 50S conventional sample | 50S ultrasonically excited sample |
|---|---|---|
| **Data collection and processing** | | |
| Microscope | Titan Krios G4i | Titan Krios G4i |
| Camera | Falcon 4i | Falcon 4i |
| Energy filter | 10 eV (SelectrisX) | 10 eV (SelectrisX) |
| Magnification | 42,000 | 42,000 |
| Voltage (kV) | 300 | 300 |
| Tilt Range (°) | +60 to -60 | +60 to -60 |
| Step (°) | 3.0 | 3.0 |
| Frames per tilt angle | 8 | 8 |
| Dose rate ($e^{-}/Å^{2}\ s^{-1}$) | 1.96 | 1.96 |
| Total dose ($e^{-}/Å^{2}$) | 120 | 120 |
| Defocus range (µm) | -5 to -6 | -5 to -6 |
| Pixel size (Å) | 3.06 | 3.06 |
| Image format | EER | EER |
| Holes imaged (no.) | 30 | 14 |

**9 | Cryo-EM data collection and processing statistics**

**Supplementary Table 2:** Data collection parameters and single particle processing statistics for the 50S ribosomal subunit.

| | 50S conventional sample (EMD-54830) | 50S ultrasonically excited sample (EMD-54829) |
|---|---|---|
| **Data collection and processing** | | |
| Microscope | Titan Krios G4i | Titan Krios G4i |
| Camera | Falcon 4i | Falcon 4i |
| Energy filter | N/A | N/A |
| Magnification | 96,000 | 96,000 |
| Voltage (kV) | 300 | 300 |
| Electron exposure ($e^{-}$/Å$^2$) | 50 | 50 |
| Defocus range (μm) | -0.4 – -1.2 | -0.4 – -1.2 |
| Pixel size (Å) | 0.830 | 0.830 |
| Image format | EER | EER |
| Symmetry imposed | $C_1$ | $C_1$ |
| Initial particle images (no.) | 238,747 | 625,311 |
| Final particle images (no.) | 50'000 | 50'000 |
| Map resolution (Å) | 2.8 | 2.8 |
| FSC threshold | 0.143 | 0.143 |
| Map resolution range (Å) | 3.18 – 2.46 | 3.06 – 2.61 |
| Map sharpening *B* factor (Å$^2$) | 29.7 | 33.0 |
| Orientation diagnostics | | |
| cFAR | 0.35 | 0.58 |
| SCF* | 0.66 | 0.83 |

**Supplementary Table 3:** Data collection parameters and single particle processing statistics for C-reactive protein.

| | C-reactive protein pentamer conventional sample (EMD-54834) | C-reactive protein pentamer ultrasonically excited sample (EMD-54833) | C-reactive protein decamer conventional sample (EMD-54836) | C-reactive protein decamer ultrasonically excited sample (EMD-54835) |
|---|---|---|---|---|
| **Data collection and processing** | | | | |
| Microscope | Titan Krios G4i | Titan Krios G4i | Titan Krios G4i | Titan Krios G4i |
| Camera | Falcon 4i | Falcon 4i | Falcon 4i | Falcon 4i |
| Energy filter | N/A | N/A | N/A | N/A |
| Magnification | 96,000 | 96,000 | 96,000 | 96,000 |
| Voltage (kV) | 300 | 300 | 300 | 300 |
| Electron exposure ($e^{-}/Å^{2}$) | 50 | 50 | 50 | 50 |
| Defocus range (µm) | -1.0 – -2.4 | -1.0 – -2.4 | -1.0 – -2.4 | -1.0 – -2.4 |
| Pixel size (Å) | 0.830 | 0.830 | 0.830 | 0.830 |
| Image format | EER | EER | EER | EER |
| Symmetry imposed | $C_5$ | $C_5$ | $C_1$ | $C_1$ |
| Initial particle images (no.) | 5,448,069 | 3,905,203 | 5,448,069 | 3,905,203 |
| Final particle images (no.) | 30’000 | 30’000 | 50’000 | 50’000 |
| Map resolution (Å) | 4.3 | 3.7 | 3.3 | 3.2 |
| FSC threshold | 0.143 | 0.143 | 0.143 | 0.143 |
| Map resolution range (Å) | 4.63 – 3.61 | 3.93 – 3.42 | 3.51 – 3.01 | 3.35 – 2.88 |
| Map sharpening *B* factor ($Å^{2}$) | 147.6 | 118.3 | 78.2 | 66.1 |
| Orientation diagnostics | | | | |
| cFAR | 0.47 | 0.64 | 0.68 | 0.67 |
| SCF* | 0.80 | 0.90 | 0.89 | 0.90 |

**Supplementary Table 4:** Data collection parameters and single particle processing statistics for hemagglutinin (jetted samples).

| | Hemagglutinin conventional sample (EMD-54832) | Hemagglutinin ultrasonically excited sample (EMD-54831) |
|---|---|---|
| **Data collection and processing** | | |
| Microscope | Titan Krios G4i | Titan Krios G4i |
| Camera | Falcon 4i | Falcon 4i |
| Energy filter | N/A | N/A |
| Magnification | 96,000 | 96,000 |
| Voltage (kV) | 300 | 300 |
| Electron exposure ($e^{-}/Å^2$) | 50 | 50 |
| Defocus range (μm) | -1.0 – -2.4 | -1.0 – -2.4 |
| Pixel size (Å) | 0.830 | 0.830 |
| Image format | EER | EER |
| Symmetry imposed | $C_3$ | $C_3$ |
| Initial particle images (no.) | 5,566,288 | 2,847,316 |
| Final particle images (no.) | 123’295 | 28’251 |
| Map resolution (Å) | 2.7 | 2.7 |
| FSC threshold | 0.143 | 0.143 |
| Map resolution range (Å) | 19.15 – 2.38 | 2.96 – 2.44 |
| Map sharpening *B* factor ($Å^2$) | 77.0 | 66.1 |
| Orientation diagnostics | | |
| cFAR | 0.00 | 0.58 |
| SCF* | 0.25 | 0.81 |

**Supplementary Table 5:** Data collection parameters and single particle processing statistics for hemagglutinin (plunged samples).

| | Hemagglutinin conventional sample (EMD-58182) | Hemagglutinin ultrasonically excited sample (EMD-58181) |
|---|---|---|
| **Data collection and processing** | | |
| Microscope | Titan Krios G4i | Titan Krios G4i |
| Camera | Falcon 4i | Falcon 4i |
| Energy filter | N/A | 10 eV |
| Magnification | 96,000 | 165,000 |
| Voltage (kV) | 300 | 300 |
| Electron exposure ($e^{-}/Å^2$) | 50 | 50 |
| Defocus range (μm) | -1.0 – -2.4 | -1.0 – -2.4 |
| Pixel size (Å) | 0.830 | 0.732 |
| Image format | EER | EER |
| Symmetry imposed | $C_3$ | $C_3$ |
| Initial particle images (no.) | 4,079,280 | 3,578,779 |
| Final particle images (no.) | 298,118 | 84,114 |
| Map resolution (Å) | 3.1 | 2.0 |
| FSC threshold | 0.143 | 0.143 |
| Map resolution range (Å) | 12.34 – 2.64 | 1.90 – 2.00 |
| Map sharpening *B* factor ($Å^2$) | 119.8 | 48.1 |
| Orientation diagnostics | | |
| cFAR | 0.01 | 0.72 |
| SCF* | 0.37 | 0.90 |

### 10 | References
